# Lineage tracing in rhabdoid tumor organoids and orthotopic xenografts reveals convergent transcriptional states

**DOI:** 10.64898/2026.11.03.761870

**Authors:** Giulia Perticari, Jeff DeMartino, Arianna Fumagalli, Dilara Ayyildiz, Sofia Doulkeridou, Lennart Kester, Jolanda Kooiman, Thanasis Margaritis, Jarno Drost

## Abstract

Malignant rhabdoid tumors (MRTs) are aggressive pediatric malignancies driven by loss of a single SWI/SNF subunit in an otherwise near-silent genome, yet they display extensive transcriptional heterogeneity. Whether this heterogeneity reflects the outgrowth of transcriptionally distinct clones, or states adopted independently of lineage, is unknown. Here we combined lentiviral lineage barcoding with single-cell RNA sequencing in two patient-derived renal MRT organoid models, followed through long-term *in vitro* culture and in a novel orthotopic xenograft setting. In both modalities, tumors and cultures became progressively restricted to a small number of lineages, and the clones that expanded were transcriptionally distinct from one another. Decomposing into gene expression programs showed that some transcriptional states were used broadly across lineages while others were confined to a few, and that lineages expanding independently, in separate cultures and separate animals, converged on the same restricted states. Individual lineages generally maintained their transcriptional identity across passaging and across anatomical sites, including in metastases, which arose from clones already expanded in the matched primary tumor. Which lineages came to dominate, however, differed between animals given the same starting population. Clonal outcome is therefore variable while the transcriptional states available to a tumor are drawn from a constrained and recurring repertoire.

## Introduction

Pediatric tumors are typically characterized by a remarkably low mutational burden compared with adult cancer^1,2^. This genetic simplicity contrasts with substantial transcriptional diversity^3^, suggesting an important contribution of non-genetic mechanisms to malignant phenotypes.

Malignant rhabdoid tumors (MRTs) represent a striking example of this apparent paradox. MRTs are rare, highly aggressive embryonal tumors that are often metastatic at diagnosis^4^. They are driven by the biallelic loss of *SMARCB1*, or in rare cases *SMARCA4*, both members of the SWI/SNF chromatin remodeling complex^5,6^. Despite this, they are characterized by a remarkable degree of heterogeneity across multiple levels. At the anatomical level, MRTs can arise intra-cranially (atypical teratoid rhabdoid tumor, ATRT) and at different extra-cranial sites, including the kidney, liver, and soft tissues (ecMRT). This heterogeneity extends to the histological level, with distinct cell morphologies and immunohistochemical profiles coexisting within MRTs^7–10^. At the molecular level, bulk epigenetic and transcriptomic analyses have identified distinct MRT subgroups^11–13^, while recent single-cell multiomic profiling has revealed subtype-specific cellular states in ATRTs^14^. How these genetically simple tumors give rise to such extensive phenotypic heterogeneity and whether this heterogeneity contributes functionally to tumor evolution are still unresolved questions.

The simple genetic landscape of MRTs makes them particularly suitable for studying non-genetic determinants of tumor evolution. Unlike other tumor entities, however, the limited number of endogenous mutations restricts their use as natural lineage markers to distinguish and track individual clones. Prospective lineage tracing using cellular barcoding technologies, such as genetic barcoding, is a powerful tool to overcome this limitation and reconstruct clonal trajectories at high resolution^15^. Previous studies have used these approaches to define the contribution of individual clones to tumor growth, metastatic dissemination and therapy resistance^16–20^. When coupled with single-cell transcriptomic profiling, lineage tracing has been used to link clonal identity to cell state^21–28^, and thus identify molecular programs associated with distinct clonal behaviors.

To study clonal evolution of primary tumor and metastasis *in vivo*, it is important to recapitulate the tissue context in which the tumor naturally develops. This can be achieved using orthotopic xenograft models, in which tumor cells are implanted into their tissue of origin. These models can also generate metastases that resemble those observed in patients^29,30^. For MRT of the kidney (MRTK), orthotopic xenograft models have been developed by renal transplantation of established cell lines^31^, which only partially recapitulate the phenotypic heterogeneity of the tumor of origin. Patient-derived MRTK organoids offer a more faithful representation of tumor heterogeneity^32^, however they have been used only in subcutaneous xenografts^33,34^. Therefore, an *in vivo* MRTK model that preserves the heterogeneity of the original tumor within its native renal environment is still lacking.

In this study, we develop a novel *in vivo* MRTK model through orthotopic xenotransplantation of patient-derived organoids into the kidneys of immunodeficient mice. The resulting tumors exhibit clinically relevant disease progression and spontaneously metastasize to distant body sites, including the lungs, brain, and lymph nodes. To resolve the clonal dynamics underlying this progression, we combine the model with a high-complexity lentiviral barcode library and single-cell RNA-sequencing (scRNA-seq), allowing us to link clonal identity and transcriptional state across organoids, primary tumors, and metastases.

## Results

### Lineage tracing reveals clonal dynamics during MRTK organoid growth

To track clonal and cell-state dynamics in MRTK cells, we adapted a lentiviral lineage-tracing system^35^ comprising a library of more than 30,000 unique 25-nucleotide barcodes positioned in the 3′ untranslated region of an EF1α promoter-driven Puromycin–PSA–H2B–mNeon fusion gene (Fig. 1a). Bulk DNA-sequencing (DNA-seq) of the integrated barcode construct enabled high-throughput quantification of population-level clonal dynamics. As each barcode was embedded within an expressed, polyadenylated transcript, clonal identity could also be captured together with transcriptional state using scRNA-seq.

**Figure 1:**
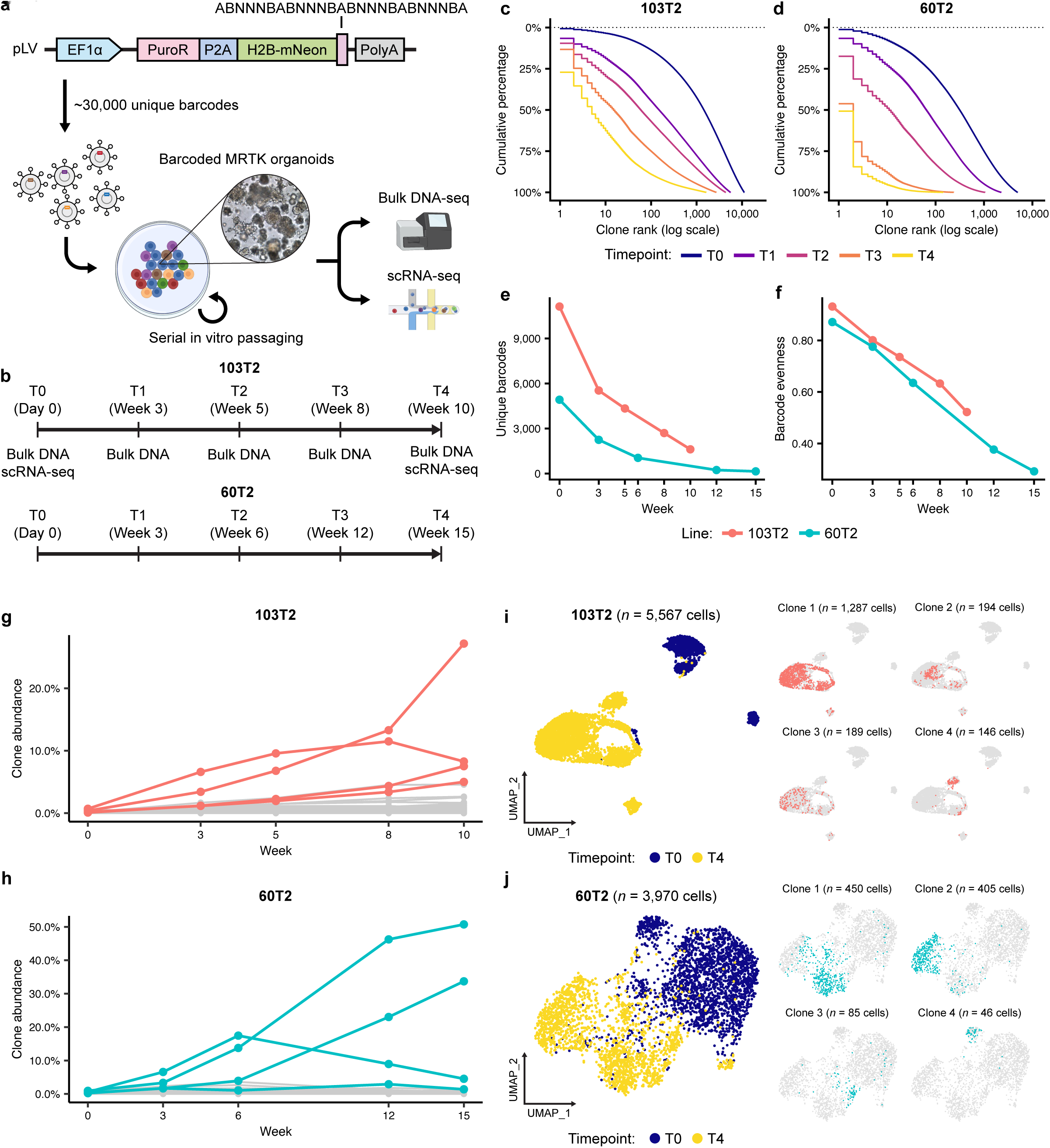
Lineage tracing of MRTK organoids during long-term culture reveals progressive clonal restriction. **(a)** Schematic of the lentiviral lineage-tracing construct and experimental workflow. **(b)** Sampling timeline. Barcode composition was measured by bulk DNA-seq at five timepoints (T0–T4); scRNA-seq was performed at T0 and T4. **(c, d)** Cumulative barcode abundance against clone rank (log scale) at each timepoint for 103T2 (c) and 60T2 (d). Colors indicate timepoint. **(e)** Number of unique barcodes recovered per timepoint in each line. **(f)** Pielou’s evenness index per timepoint in each line. **(g, h)** Relative abundance of individual barcodes across the time course in 103T2 (g) and 60T2 (h). Colored lines indicate clones highlighted in (i, j); all other clones are shown in gray. **(i, j)** UMAP projections of scRNA-seq data for 103T2 (i, n = 5,567 cells) and 60T2 (j, n = 3,970 cells). Left, cells colored by timepoint. Right, the four most abundant T4 lineages highlighted individually, with cell numbers indicated; all other cells are shown in gray.

To characterize clonal dynamics during long-term *in vitro* culture, we independently transduced two previously established patient-derived MRTK organoid lines, 103T2 and 60T2^32^. Barcoded organoids were then passaged weekly (at a ratio of 1:4) for up to 15 weeks, during which barcode composition was quantified by bulk DNA-seq at five timepoints spanning the culture period (T0–T4; Fig. 1b-d). Immediately following antibiotic selection (T0), 11,125 and 4,921 unique barcodes were recovered from 103T2 and 60T2, respectively (Fig. 1c-e). Importantly, barcode evenness (Pielou’s Evenness Index) was relatively high in both lines (0.931 and 0.871), indicating that neither population was dominated by a small number of clones at the outset of the experiment (Fig. 1f).

Clonal complexity declined markedly with continued passaging in both lines. The number of unique barcodes recovered fell to 1,617 at Week 10 (T4) in 103T2, and 148 at Week 15 (T4) in 60T2 (Fig. 1e), reductions of approximately 7- and 33-fold respectively. This loss was accompanied by a progressive redistribution of the population towards a small number of lineages. At T0, 1,450 clones accounted for half of all reads in 103T2 and 314 in 60T2. By T4, the same fraction was contributed by only 4 clones in 103T2 and a single clone in 60T2 (Fig. 1c, d). Evenness declined in parallel to 0.522 in 103T2 and 0.292 in 60T2 (Fig. 1f). The surviving populations therefore did not simply lose rare clones at random, but became increasingly unequal in clone size.

To identify the lineages responsible, we tracked individual barcodes across the time course. Compared with T0, only 80 (103T2) and 16 (60T2) barcodes increased more than four-fold in relative abundance by T4, while the large majority of lineages declined (Fig. 1g, h). Despite their small number, these expanding lineages came to constitute 88.9% and 96.5% of the final population, compared with 4.01% and 2.64% at T0 (Fig. 1g, h; Supplementary Fig. S1a, b). Such progressive, directional expansion is consistent with selection acting on pre-existing clonal differences. We note, however, that serial passaging also imposes repeated population bottlenecks, and drift alone can reduce diversity and skew clone sizes in the absence of any fitness difference, consistent with a smaller set of lineages that rose at intermediate passages before contracting again by T4 (Fig. 1g,h).

To relate clonal identity to transcriptional state, we performed scRNA-seq on barcoded organoids at T0 and T4 in both lines (Fig. 1b). After quality control filtering, we obtained 5,567 and 3,970 cells for 103T2 and 60T2 respectively, of which 88.9% and 65.5% carried a detectable barcode (Supplementary Fig. S1c-f). To establish that clonal structure could be read directly from single-cell data, we compared barcode frequencies measured by bulk DNA-seq with those obtained from scRNA-seq of the same samples (Supplementary Fig. S1g). At T4, agreement was high in both lines (Spearman ρ = 0.58 and 0.9), and barcodes detected in scRNA-seq accounted for 89.2% and 96.2% of total clonal abundance in the corresponding bulk libraries, indicating that single-cell profiling captured essentially all of the dominant clonal mass. Agreement was lower at T0, particularly in 103T2 (ρ = 0.17), but this reflects the near-uniform frequency distribution at that timepoint: with several thousand clones present at comparable abundance, there is little true variation for the two assays to agree upon, and the correlation is sensitive to sampling noise in both. Consistent with this, the less even 60T2 T0 sample gave a notably higher correlation (ρ = 0.56). Taken together, these comparisons indicate that scRNA-seq recovers the dominant clonal structure of these populations, supporting its use to link lineage identity with transcriptional state.

Finally, uniform manifold approximation and projection (UMAP) of the scRNA-seq data showed cells from T0 and T4 occupying largely non-overlapping regions of the embedding in both lines, consistent with a substantial shift in transcriptional state over the course of passaging (Fig. 1i, j). Highlighting the four most abundant T4 lineages revealed that these were not uniformly distributed across this space, with individual clones concentrated in distinct regions of the endpoint population. Thus, lineages expanding during passaging occupy distinct transcriptional states.

### Distinct transcriptional states underlie clonal expansion in MRT organoids

To characterize the transcriptional states occupied by the lineages that expanded *in vitro*, we used non-negative matrix factorization (NMF) to decompose the scRNA-seq data into coherent gene expression programs (GEPs). In 103T2, this yielded six programs, which we annotated from their top-weighted genes and the biological processes enriched among them (Fig. 2a; Supplementary Fig. S2a,b). These comprised a stress and immediate-early response program (Stress/AP-1; *FOSL1*, *ATF3*, *EGFR*); a proliferative program of cell-cycle and mitotic spindle genes (Cell cycle; *MKI67*, *TOP2A*, *CCNB1*); a program distinguished by axon guidance and neurotrophic signaling expression (Neuronal; *NTN1*, *NRG3*, *EFNA5*); a program enriched for developmental transcription factors and receptors (Developmental; *RUNX2*, *FGFR1*, *EMX2*); a program of cell-surface and matrix-remodeling genes (Mesenchymal; *THY1*, *CD44*, *LOXL2*); and a program defined by neurodevelopmental genes (*NLGN1*, *ROBO1*, *RELN*), which we refer to as “Progenitor-like”.

**Figure 2:**
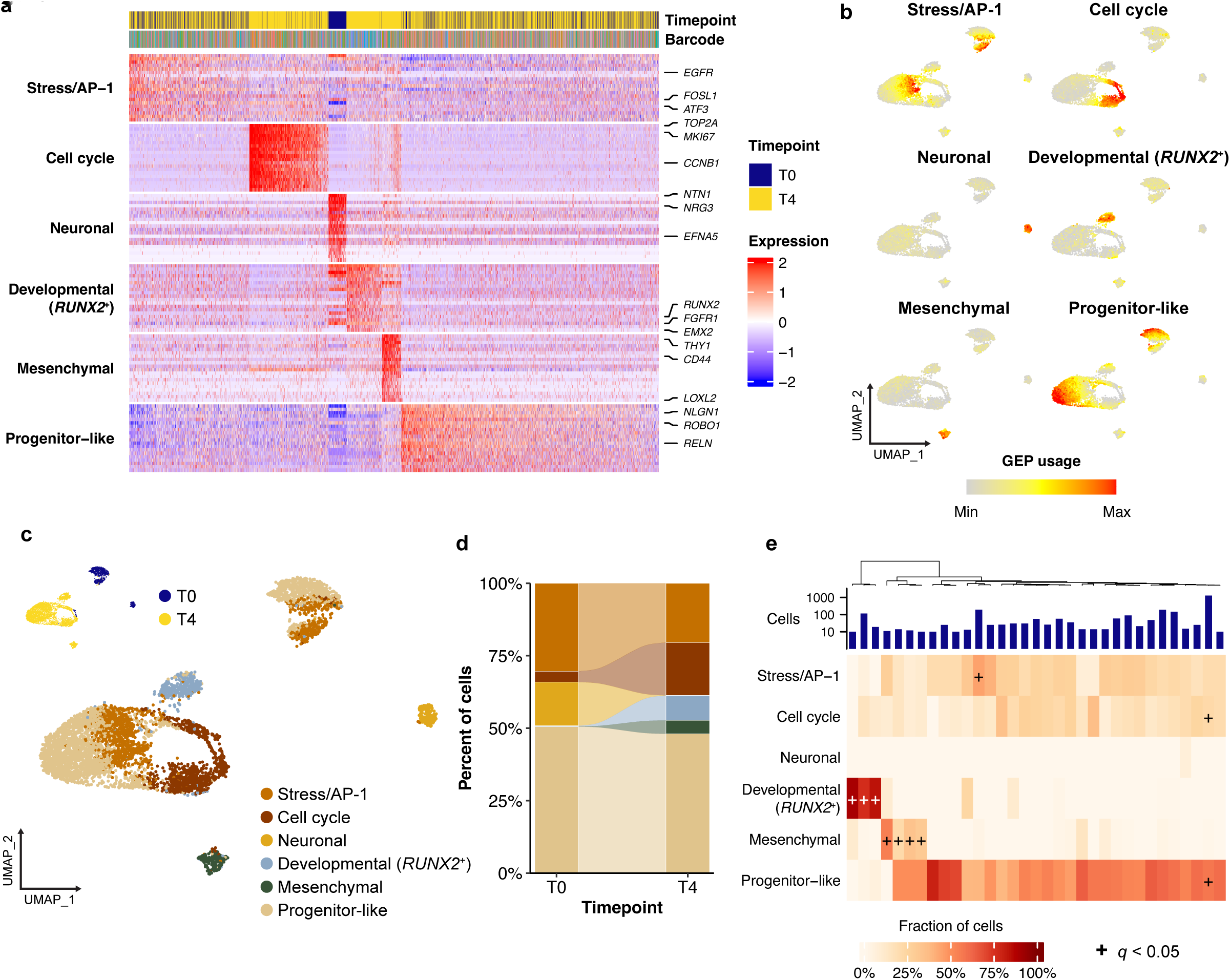
Analysis of transcriptional programs in 103T2 organoids show that loss of clonal diversity is accompanied by the emergence of discrete cell states. **(a)** Scaled expression of the top-weighted genes for each of six NMF-derived gene expression programs (GEPs). Rows are grouped by program; columns are cells, ordered by assigned program and by usage of that program. Annotation tracks indicate timepoint and barcode identity. Selected marker genes are labeled at right. **(b)** UMAP projections colored by per-cell usage of each program. **(c)** UMAP projection colored by the program with the highest usage in each cell. Inset, the same embedding colored by timepoint. **(d)** Proportion of cells assigned to each program at T0 and T4. Bands connect the same program between timepoints. **(e)** Fraction of each lineage’s cells assigned to each program, for the 33 lineages represented by at least 10 cells at T4. Columns are lineages, ordered by hierarchical clustering of program composition; the barplot above shows the number of cells per lineage (log scale). Plus signs mark program-lineage combinations significantly enriched relative to the remaining cells of the same sample (hypergeometric test, Benjamini-Hochberg adjusted q < 0.05).

Programs differed in how broadly they were used (Fig. 2b; Supplementary Fig. S2b). The neuronal, developmental and mesenchymal programs were each restricted to smaller, discrete populations of cells, whereas the stress/AP-1 and progenitor-like programs were used at moderate levels across most of the population. Assigning each cell to its most-used program reflected this structure, with the restricted programs marking well-separated groups and the broadly used programs accounting for the bulk of cells (Fig. 2c).

Program composition changed substantially between T0 and T4 (Fig. 2d). The T0 population was dominated by the progenitor-like and stress/AP-1 programs, which together accounted for 80.9% of cells, with a further 15.1% assigned to the neuronal program. By T4 the proliferative program had expanded from 3.8% to 18.2% of cells, and the developmental and mesenchymal programs, essentially absent at T0 (0.24% and 0.08%), together accounted for 13.2% of cells. Conversely, the neuronal program was almost entirely lost, falling to 0.05% of cells. Notably, the programs that emerged over passaging were those confined to discrete subsets of cells, whereas the broadly used programs changed comparatively little. The endpoint population was therefore composed of fewer clonal lineages than at T0, but was partitioned into a greater number of distinct transcriptional states, indicating that the loss of clonal diversity was accompanied not by transcriptional homogenization, but by the emergence of discrete cell states within a restricted set of clones.

We next asked whether these transcriptional states were associated with clonal identity. Restricting the analysis to the 33 lineages represented by at least 10 cells at T4, we determined the fraction of each clone’s cells assigned to each program and tested for enrichment against the remaining cells of the same sample (Fig. 2e). Programs differed markedly in the extent to which they were clone-associated. The progenitor-like and proliferative programs were used at comparable levels by nearly all lineages, consistent with their interpretation as more general states. In contrast, the developmental program was almost entirely restricted to 3 clones, in each of which it accounted for more than 76% of cells (q < 0.05), while being negligible elsewhere. A further 4 clones were significantly enriched for the mesenchymal program. Clustering lineages by their program composition grouped these clones together, indicating that independent lineages converged on a set of shared transcriptional states, rather than each adopting a private one.

We next applied the same approach to the scRNA-seq data from 60T2, which had undergone a more severe clonal restriction. NMF decomposition again yielded six GEPs (Fig. 3a; Supplementary Fig. S3a,b): a proliferative program (Cell cycle; *TOP2A*, *MKI67*, *CCNB1*); a program marked by matrix metalloproteinases and collagens (ECM/invasion; *MMP3*, *COL12A1*, *FGF1*); a WNT and developmental program (WNT/developmental; *WNT5A*, *SIX1*, *FZD7*); a program marked by hypoxia-response and sterol biosynthesis genes (Hypoxia/sterol; *LOX*, *SQLE*, *PLOD2*); a neuronal and glial program (Glial-like; *PDGFRA*, *CLU*, *GRIA2*); and, as in 103T2, a broadly used program marked by neuronally expressed, developmentally related genes (Progenitor-like; *AUTS2*, *BICC1*).

**Figure 3:**
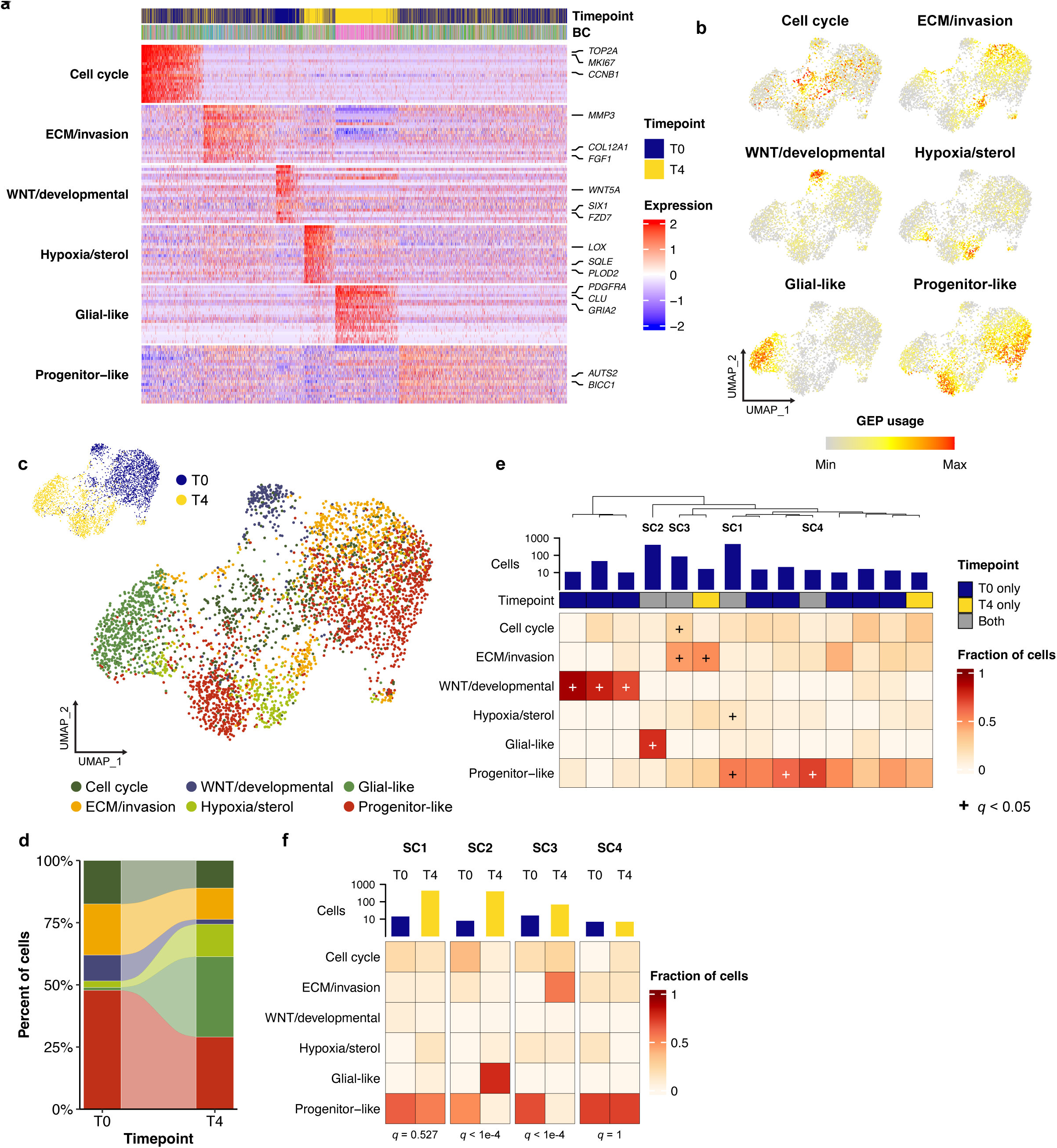
Analysis of transcriptional programs in 60T2 organoids reveals that individual lineages differ in and change their program composition over passaging. **(a)** Scaled expression of the top-weighted genes for each of six NMF-derived gene expression programs (GEPs). Rows are grouped by program; columns are cells, ordered by assigned program and by usage of that program. Annotation tracks indicate timepoint and barcode identity. Selected marker genes are labeled at right. **(b)** UMAP projections colored by per-cell usage of each program. **(c)** UMAP projection colored by the program with the highest usage in each cell. Inset, the same embedding colored by timepoint. **(d)** Proportion of cells assigned to each program at T0 and T4. Bands connect the same program between timepoints. **(e)** Fraction of each lineage’s cells assigned to each program, for the 14 lineages represented by at least 10 cells. Annotation tracks indicate the number of cells per lineage (log scale) and whether the lineage was detected at T0 only, T4 only, or both. Lineages compared across timepoints in (f) are labeled SC1–SC4. Plus signs as in Figure 2e. **(f)** Program composition of the four lineages detected at both T0 and T4, shown separately for each timepoint. Barplots indicate cells per lineage per timepoint (log scale). Fisher’s exact test *q*-values for the difference in composition between timepoints are given below each pair.

Previous work mapping MRTK along neural crest differentiation placed 103T2 and 60T2 at distinct points along this trajectory^36^. Consistent with this, only two programs corresponded directly between the two lines (Supplementary Fig. S3c): the proliferative programs, as expected for a state defined by cell-cycle genes, and the progenitor-like programs, whose most highly weighted genes overlapped substantially between the two decompositions. The remaining programs were specific to one or the other model, in keeping with the different developmental positions of the two tumors of origin.

As in 103T2, program composition shifted markedly over passaging (Fig. 3c, d). At T0 the population was dominated by the ECM/invasion and progenitor-like programs (20.6% and 47.8% of cells), with the proliferative and WNT/developmental programs accounting for a further 17.5% and 10.4%. By T4 the glial-like program had expanded from 1.2% to 32.3% of cells and the hypoxia/sterol program from 2.6% to 13.1%, while the ECM/invasion, proliferative and WNT/developmental programs all declined (12.6%, 11.1%, and 1.9% respectively).

We next examined whether program usage was associated with clonal identity, as it was in 103T2. Across the 14 lineages represented by at least 10 cells, programs again differed in the extent to which they were clone-restricted (Fig. 3e). The proliferative and progenitor-like programs were used at broadly comparable levels across most lineages, whereas the WNT/developmental and glial-like programs were each significantly enriched in a small number of clones. These restricted programs differed in when the lineages using them were sampled: the clones enriched for the WNT/developmental program were detected only at T0, consistent with the loss of this program from the population over passaging, whereas the glial-like program was enriched in only a single lineage, despite accounting for a substantial share of cells at T4. Clustering lineages by their program composition grouped clones with similar usage together, showing, again, that independently arising lineages converged on the same transcriptional states.

Notably, we identified 4 clones represented by at least 10 cells at both T0 and T4 (SC1–SC4), enabling us to directly compare program composition between the two timepoints within individual lineages (Fig. 3e, f). While SC1 and SC4 retained a predominantly progenitor-like composition (q = 0.527 and q = 1; Fisher’s exact test), SC2 and SC3 shifted substantially towards the glial-like and ECM/invasion programs respectively (both q < 1 × 10⁻⁴). That the same lineage could change its program composition over passaging suggests that transcriptional state is plastic within a lineage, and that at least part of the shift observed at the population level reflects changes within persisting lineages, rather than the replacement of one set of clones by another. At the same time, the extent of change differed markedly between lineages, and clonal identity remained associated with program usage at the endpoint (Fig. 3e). Clonal identity and transcriptional state are therefore coupled, but not deterministically.

Taken together, we show for the first time that MRTK cells undergo clonal evolution in culture: prolonged passaging drove a substantial loss of clonal diversity in both lines, accompanied by a reorganization of transcriptional state. Given the genomic stability of MRTs^37^, and of these organoid models in extended culture^32^, this is unlikely to reflect the acquisition of new genetic drivers. The lineages that persisted were not simply those in the most proliferative state: in both lines the proliferative program was used at comparable levels across nearly all lineages, and in 60T2 its share of the total population declined over passaging. What distinguished surviving clones from one another were instead the restricted, non-cycling programs. Nor was the reorganization only a matter of which clones survived: a single lineage could span several transcriptional states at the endpoint, and lineages tracked from T0 to T4 sometimes differed in their program composition. The shift is therefore driven by plasticity within persisting clones as well as by selection between them. Although the programs themselves differed between the two lines, this reorganization of transcriptional state was common to both.

### Lineage tracing in orthotopic MRTK xenografts reveals extensive clonal dynamics *in vivo*

Having characterized clonal and transcriptional dynamics under *in vitro* culture conditions, we next asked how the same lineages behave *in vivo*, where cells face additional selective pressures during engraftment, tumor growth and dissemination. To this end, we established an orthotopic xenotransplantation model in which barcoded MRTK organoids were injected under the renal capsule of immunodeficient mice, generating tumors that histologically recapitulated the disease in patients and maintained loss of SMARCB1/INI1 expression (Fig. 4a-c). Importantly, tumors spontaneously metastasized to distant organs, including the lung, liver, brain and lymph nodes, mirroring the metastatic sites observed in patients^4^ (Fig. 4b).

**Figure 4:**
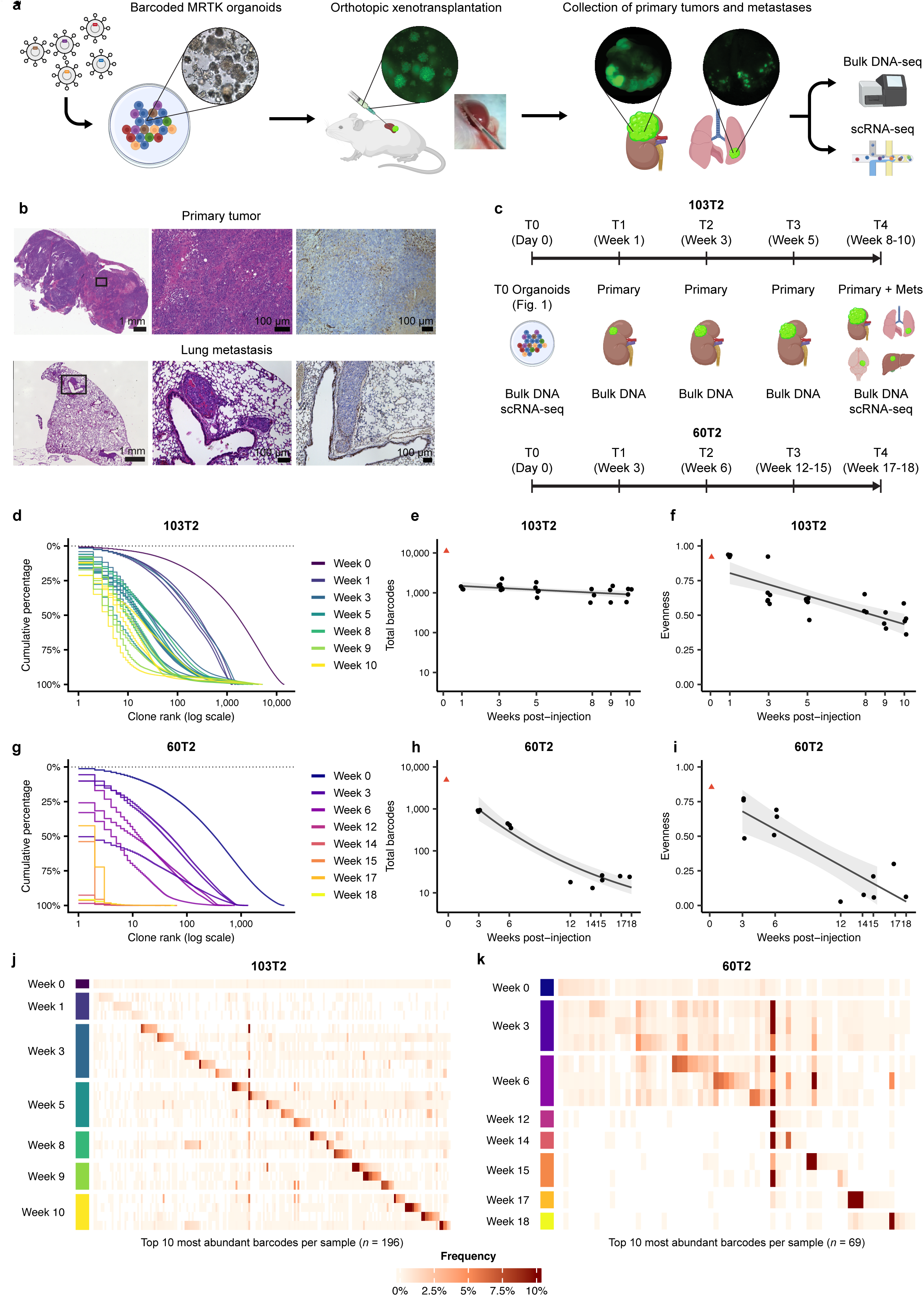
Clonal dynamics of MRTK orthotopic xenografts reavel engraftment-associated bottleneck and progressive clonal dominance during tumor progression. **(a)** Schematic of the orthotopic xenotransplantation workflow. Barcoded organoids were injected under the renal capsule of NSG mice, and primary tumors and metastases were collected for bulk DNA-seq and scRNA-seq. **(b)** Representative histology of a primary tumor (top row) and a lung metastasis (bottom row). Left and center, hematoxylin and eosin; right, INI1 immunohistochemistry. Scale bars, 1 mm (left) and 100 µm (center, right). **(c)** Sampling timelines for 103T2 and 60T2, indicating the tissue collected and assay performed at each timepoint. **(d, g)** Cumulative barcode abundance against clone rank (log scale) for individual tumors, colored by week collected, post-injection, in 103T2 (d) and 60T2 (g). **(e, h)** Number of barcodes recovered per tumor against week collected, post-injection in 103T2 (e) and 60T2 (h). Each point is one tumor; the red triangle indicates the injected organoid pool (T0). Line and shaded band show glm with 95% confidence interval; the T0 sample was excluded from the fit. **(f, i)** Pielou’s evenness index per tumor, plotted as in (e, h). Line and shaded band show lm with 95% confidence interval; the T0 sample was excluded from the fit. **(j, k)** Frequency of the union of the ten most abundant barcodes per sample across all tumors, for 103T2 (j, *n* = 196 barcodes) and 60T2 (k, *n* = 69 barcodes). Rows are individual tumors grouped by week; columns are barcodes, ordered by the sample in which each reaches its highest frequency. White indicates barcodes not detected.

Both barcoded lines were injected into cohorts of 25 (103T2) or 21 (60T2) mice, and animals were sacrificed at a series of timepoints spanning tumor development: weeks 1 to 10 for 103T2, and weeks 3 to 18 for 60T2, reflecting the slower growth of the latter (Fig. 4c; Supplementary Table S1). Tumor growth was monitored throughout by bioluminescence imaging (BLI; Supplementary Fig. S4a,b). At each timepoint, primary tumors from at least three mice were collected and their barcode composition determined by bulk DNA-seq. At the final timepoint, metastases were collected in addition to primary tumors for scRNA-seq. Because tumors were harvested at sacrifice, each mouse contributes a single observation and the resulting series is cross-sectional rather than longitudinal; all tumors nevertheless derive from the same barcoded organoid pool (T0), allowing clonal composition to be compared both across time and between animals.

Clonal complexity was sharply reduced during engraftment and early outgrowth *in vivo*. In the earliest tumors profiled, barcode numbers had fallen from 11,125 (103T2) and 4,921 (60T2) in the injected pools (T0 organoids) to medians of 1,245 and 918 respectively (Fig. 4e, h). Evenness, by contrast, was largely unchanged in both lines over the same interval (0.93 at T0 versus a median of 0.92 for 103T2, and 0.85 versus 0.77 for 60T2; Fig. 4f, i). The loss of complexity at engraftment was therefore not accompanied by a shift towards dominance by particular lineages, as expected if lineages were lost by chance rather than filtered by fitness, with only a small fraction of injected lineages succeeding in establishing tumors. Consistent with this, the lineages that had expanded during *in vitro* passaging were largely absent from the dominant clones of any xenograft, and vice versa (Supplementary Fig. S4c, d).

Beyond this initial bottleneck, the two lines followed different trajectories (Fig. 4d-i). In 60T2, complexity continued to decline throughout tumor development, to a median of 24 barcodes detected in tumors collected after week 15 (T4), with evenness falling in parallel to below 0.18 (Fig. 4h, i). In 103T2, by contrast, the number of barcodes recovered did not decline substantially after the first week (Fig. 4e), while evenness continued to fall (Fig. 4f). That diversity was maintained while the distribution became progressively more unequal indicates that the later phase of tumor growth in this line involved the expansion of particular lineages within a persisting pool, rather than the continued loss of clones. The two models therefore differ not only in the extent of clonal restriction but in when it occurs, with 60T2 undergoing sustained attrition and 103T2 reaching a stable complexity that is then progressively dominated by a subset of clones. The timing of clonal restriction in 60T2 mirrored the growth kinetics of the tumors. BLI remained near baseline for the first six weeks and rose steeply from week 9 onwards, over the same interval in which evenness fell dramatically, so that the emergence of clonal dominance accompanied the transition from tumor persistence to outgrowth (Supplementary Fig. S4b).

The extent to which the same lineages dominated in different animals differed between the two models. In 103T2, each tumor was dominated by a largely distinct set of clones (Fig. 4j). Among the ten most abundant barcodes in every sample, few reached comparable abundance in more than one animal, irrespective of timepoint (Fig. 4j). In 60T2, tumors harvested at earlier timepoints shared much of their abundant clonal repertoire, and these shared lineages were largely those that had been most abundant in the injected organoid pool (Fig. 4k). The latest tumors, by contrast, were each dominated by one or a small number of clones that were generally not abundant in other animals (Fig. 4k), and the transition between these patterns coincided with the sharp decline in clonal complexity in this line (Fig. 4h, i). Dominance *in vivo* was therefore established largely independently in each animal in 103T2, whereas in 60T2 the early expansion of well-represented founder lineages preceded the emergence of mainly tumor-specific clones.

### Clonal and transcriptional structure of MRTK xenografts and metastases

We next analyzed the scRNA-seq data from primary tumors at the experimental endpoint and matched metastases, allowing clonal and transcriptional relationships between primary and metastatic disease to be examined directly. Beginning with 103T2, four tumor-bearing mice were analyzed (103_1, 103_2, 103_3 and 103_4), yielding 24,780 cells in total (Fig. 5a). Metastases were collected from lung, brain, diaphragm and abdomen, giving between one and four metastatic samples per animal. Because sample sizes varied considerably, ranging from 1 to 5,169 barcoded cells, we restricted clone-level analyses to samples with at least 20 cells (Supplementary Fig. S5a). UMAP projection of the combined dataset showed that both mouse of origin and anatomical site contributed to transcriptional differences between cells, although neither fully accounted for the observed structure, with cells from different animals, particularly from primary tumors, occupying overlapping regions of the embedding (Fig. 5a).

**Figure 5:**
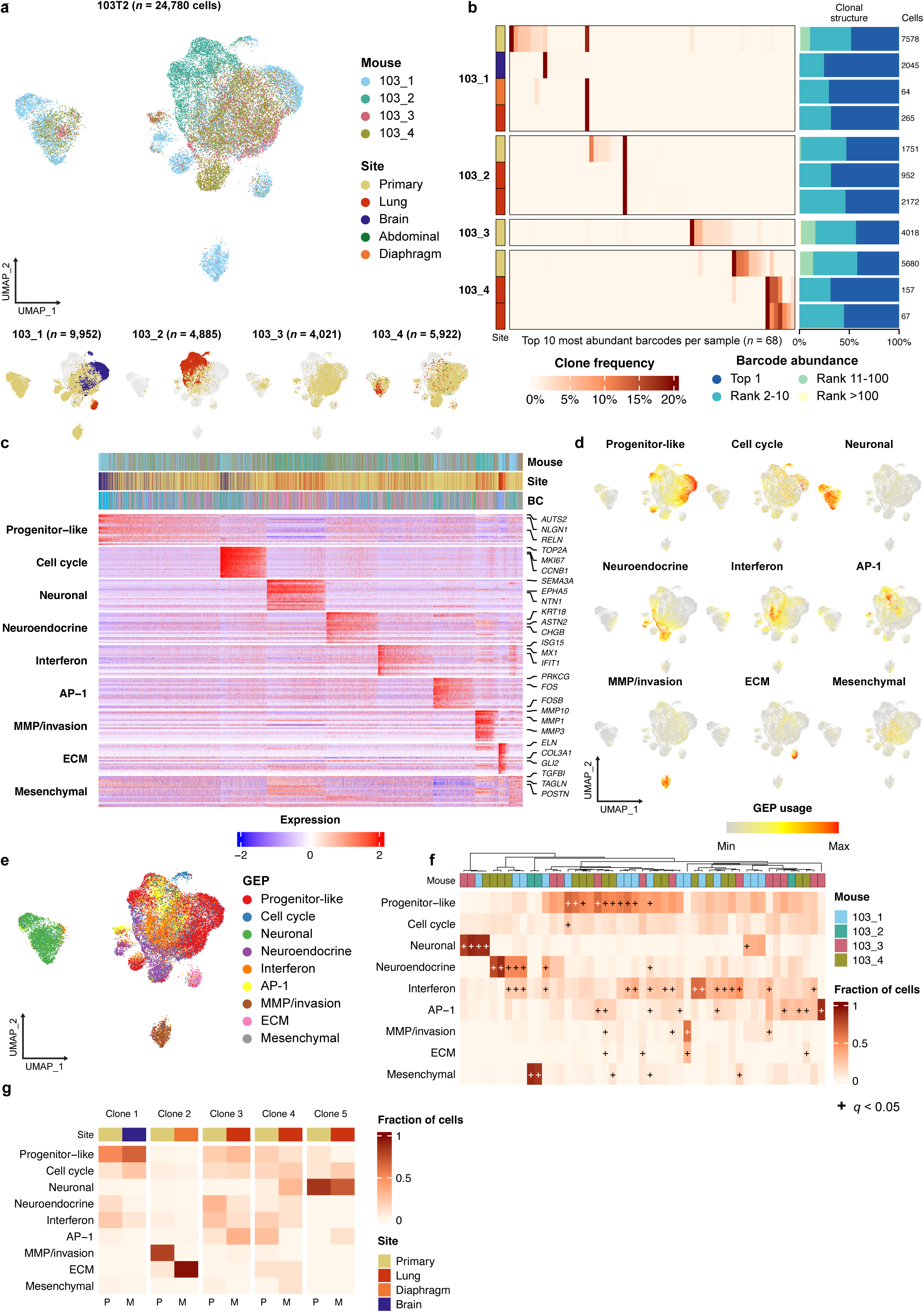
In 103T2 xenografts, metastases arise from locally expanded clones, and independently expanded lineages converge on a shared repertoir of transcriptional programs. **(a)** UMAP projection of scRNA-seq data from four 103T2-bearing mice (*n* = 24,780 cells), colored by mouse of origin. Below, cells from each mouse shown separately and colored by anatomical site, with all other cells in gray. Cell numbers per mouse are indicated. **(b)** Frequency of the union of the ten most abundant barcodes per sample (*n* = 68 barcodes). Rows are samples grouped by mouse; the left annotation indicates anatomical site. Right, the proportion of barcoded cells accounted for by the most abundant barcode and by barcodes of decreasing rank, with the total number of barcoded cells per sample given alongside. **(c)** Scaled expression of the top-weighted genes for each of nine NMF-derived programs, with annotation tracks for mouse, site and barcode. Selected marker genes are labeled at right. **(d)** UMAP projections colored by per-cell usage of each program. **(e)** UMAP projection colored by the program with the highest usage in each cell. **(f)** Fraction of each lineage’s cells assigned to each program, for lineages represented by at least 20 cells. Columns are lineages ordered by hierarchical clustering of program composition, annotated by mouse of origin. Plus signs mark significant enrichment (hypergeometric test, q < 0.05). **(g)** Program composition of the five lineages represented by at least 20 cells in both a primary tumor and a matched metastasis, shown separately for the primary (P) and metastatic (M) compartment. Site annotation indicates the anatomical origin of each sample.

Clonal composition was strongly structured by mouse. Among the ten most abundant barcodes in each sample, few clones were shared between animals, consistent with the private dominance observed by bulk DNA-seq (Fig. 5b). Within each mouse, by contrast, primary tumors and their metastases shared dominant lineages, with the exception of mouse 103_4, where both lung metastases were dominated by a shared clone that was only modestly represented in the matched primary. Metastases were also consistently more clonal than their matched primaries, with the single most abundant barcode accounting for a median of 97.9% of barcoded cells compared with 39.1% in primaries (Fig. 5b), indicating a further restriction of clonal diversity at dissemination. That metastases were dominated by the same lineages as their primaries suggests that dissemination in this model draws from clones that have already expanded locally, rather than from a distinct subpopulation with specific metastatic capacity.

NMF decomposition yielded nine transcriptional programs (Fig. 5c, d; Supplementary Fig. S5b, c): progenitor-like (*AUTS2*, *NLGN1*, *RELN*), proliferative program (Cell cycle; *TOP2A*, *MKI67*, *CCNB1*), neuronal (*SEMA3A*, *EPHA5*, *NTN1*), neuroendocrine (*KRT18*, *ASTN2*, *CHGB*), interferon (*ISG15*, *MX1*, *IFIT1*), AP-1 (*FOS*, *FOSB*, *PRKCG*), MMP/invasion (*MMP1*, *MMP3*, *MMP10*), ECM (*ELN*, *COL3A1*, *GLI2*) and mesenchymal (*TGFBI*, *TAGLN*, *POSTN*). Notably, several of these programs corresponded closely to those identified in 103T2 organoids (Supplementary Fig. S5d), indicating that cell state heterogeneity is at least partially conserved between modalities.

Program composition varied substantially between samples, but did not correspond to anatomical site (Fig. 5e, Supplementary Fig. S5e). In mouse 103_1, for example, a diaphragm and a lung metastasis were near-identical in composition, whereas the two lung metastases from mouse 103_4 differed substantially from one another. Nor was any program consistently enriched or depleted in metastases relative to their matched primary tumors. Instead, all nine programs were represented across multiple mice and sites, indicating that lineages expanding independently in different animals converge on a shared repertoire of transcriptional states.

Like in the organoid cultures, programs differed markedly in the extent to which they were clone-associated (Fig. 5f). The progenitor-like and proliferative programs were used at comparable levels across a broad swath of lineages, while the neuronal, neuroendocrine and mesenchymal programs were each restricted to a small number of lineages, in which they accounted for the majority of cells. Clustering lineages by their program composition grouped clones from different animals together, indicating that clones that arose and expanded in separate hosts converged on the same restricted transcriptional states.

Pairing primary tumors and metastases from the same mouse allowed us to compare transcriptional state within a single lineage across anatomical sites. In 103T2, five clones were represented by at least 20 cells in both a primary tumor and a matched metastasis (Fig. 5g). In four of these lineages, program composition was broadly similar in the primary tumor and the matched metastasis, differing only in the relative use of individual programs. Clone 2 was the clear exception: this lineage was dominated by the MMP/invasion program in the primary tumor and almost entirely by the ECM program in the matched metastasis. Metastatic dissemination can therefore proceed either with or without a substantial change in the transcriptional state of the disseminating lineage.

We applied the same analysis to 60T2, in which bulk barcode sequencing showed sustained clonal loss throughout tumor progression (Fig. 4h, i), profiling primary tumors and matched metastases in four mice (60_1, 60_2, 60_3 and 60_4) from which we recovered a total of 11,340 cells (Fig. 6a). Metastases were collected from the liver, spleen, and lymph nodes, as well as a lesion associated with the hepatic vasculature, resulting in one to two metastatic samples per animal. As in 103T2, sample sizes varied considerably, from 22 to 1,400 barcoded cells, and clone-level analyses were restricted to samples with at least 20 cells. UMAP projection of the combined dataset showed cells separating largely by mouse of origin, with each animal contributing one or more distinct populations and comparatively little overlap between them (Fig. 6a). This contrasts with 103T2, where cells from different animals occupied substantially overlapping regions.

**Figure 6:**
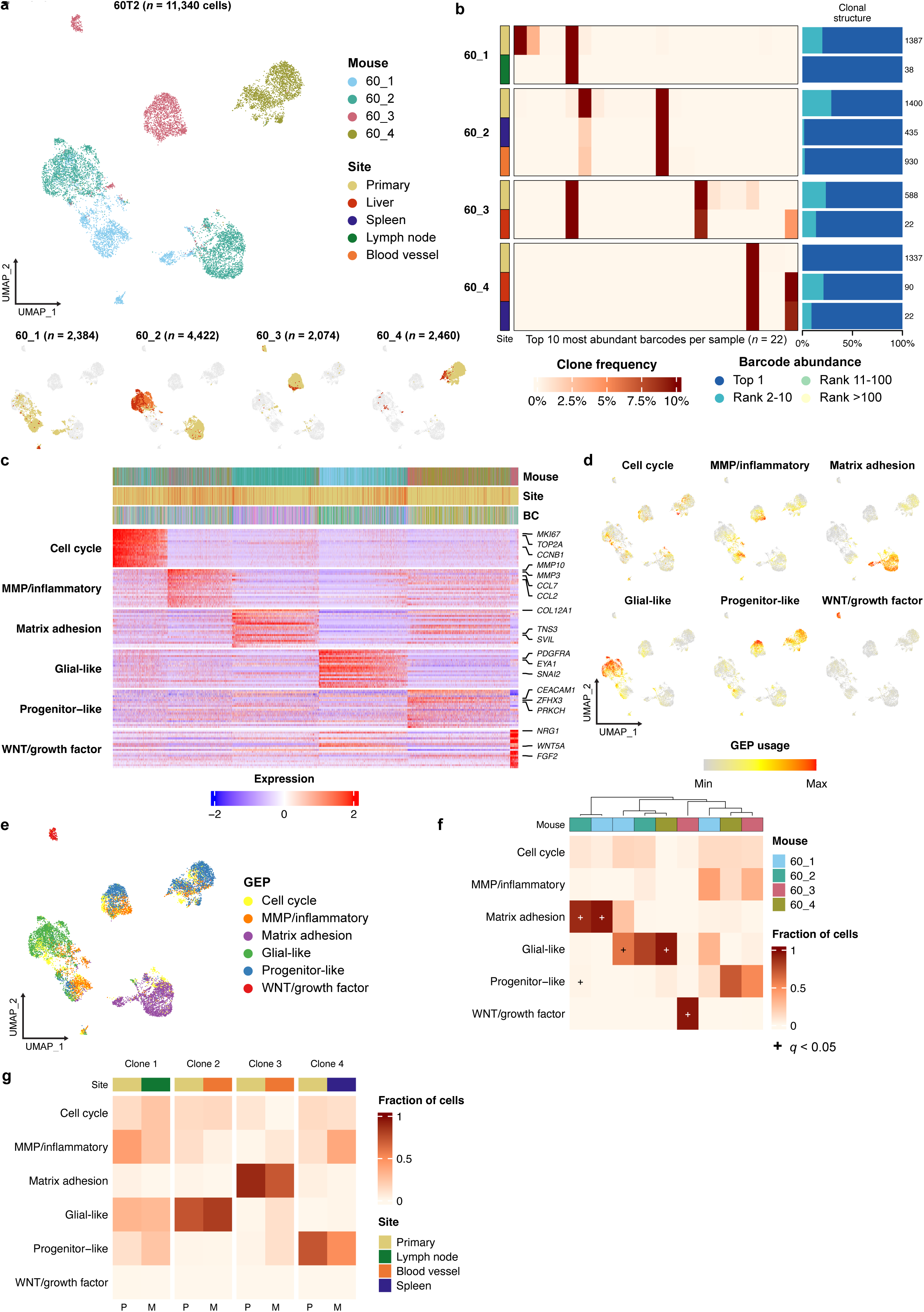
Transcriptional programs in 60T2 xenografts are clone-restricted and retained between primary tumors and metastases. **(a)** UMAP projection of scRNA-seq data from four 60T2-bearing mice (*n* = 11,340 cells), colored by mouse of origin. Below, cells from each mouse shown separately and colored by anatomical site, with all other cells in gray. Cell numbers per mouse are indicated. **(b)** Frequency of the union of the ten most abundant barcodes per sample (*n* = 22 barcodes). Rows are samples grouped by mouse; the left annotation indicates anatomical site. Right, the proportion of barcoded cells accounted for by the most abundant barcode and by barcodes of decreasing rank, with the total number of barcoded cells per sample given alongside. **(c)** Scaled expression of the top-weighted genes for each of six NMF-derived programs, with annotation tracks for mouse, site and barcode. Selected marker genes are labeled at right. **(d)** UMAP projections colored by per-cell usage of each program. **(e)** UMAP projection colored by the program with the highest usage in each cell. **(f)** Fraction of each lineage’s cells assigned to each program, for lineages represented by at least 20 cells. Columns are lineages ordered by hierarchical clustering of program composition, annotated by mouse of origin. Plus signs mark significant enrichment (hypergeometric test, *q* < 0.05). **(g)** Program composition of the four lineages represented by at least 20 cells in both a primary tumor and a matched metastasis, shown separately for the primary (P) and metastatic (M) compartment. Site annotation indicates the anatomical origin of each sample.

Clonal composition in this line was far more restricted than in 103T2. In most samples a single barcode accounted for the majority of barcoded cells, and the ten most abundant barcodes typically comprised nearly the entire population (Fig. 6b). Metastases were again somewhat more clonal than their matched primary tumors, although the difference was less pronounced than in 103T2, as primary tumors were themselves already close to monoclonal at the final timepoint. As in 103T2, dominant clones were largely private to each animal and shared between a primary tumor and its metastases, with the exception of one lineage that was dominant in both 60_1 and 60_3.

NMF decomposition yielded six GEPs (Fig. 6c, d; Supplementary Fig. S6b, c): a proliferative program (Cell cycle; *MKI67*, *TOP2A*, *CCNB1*); an MMP and inflammatory program combining matrix metalloproteinases with chemokines (*MMP3*, *MMP10*, *CCL2*, *CCL7*); an matrix adhesion program (*COL12A1*, *TNS3*, *SVIL*); a glial-like program (*PDGFRA*, *EYA1*, *SNAI2*); a WNT and growth factor program (*WNT5A*, *NRG1*, *FGF2*); and a progenitor-like program (*CEACAM1*, *ZFHX3*, *PRKCH*). As in 103T2, several of these programs corresponded closely to those identified in 60T2 organoids (Supplementary Fig. S6d), again indicating that cell state heterogeneity is at least partially conserved between modalities.

Program composition varied substantially between samples, including among the primary tumors (Fig. 6e; Supplementary Fig. S6e). Some of this variation reflected programs largely confined to a single animal, most clearly the WNT/growth factor program in 60_3. Comparing compartments within each animal, matrix adhesion usage tended to be lower and glial-like usage higher in metastases than in their matched primary tumors, in all four mice.

Relating program usage to clonal identity showed that individual lineages were strongly associated with particular programs (Fig. 6f). Most clones were dominated by a single program, and each of the restricted programs (matrix adhesion, glial-like and WNT/growth factor) accounted for the majority of cells in one or a few lineages while being negligible in the rest. As in 103T2, clustering lineages by their program composition grouped clones from different animals together rather than by mouse of origin, indicating that independently expanded lineages again converged on the same transcriptional states. One lineage was dominant in two animals (60_1 and 60_3), allowing the contribution of lineage identity to be assessed directly: these two populations were no more similar in program composition than either was to unrelated clones from other mice. Transcriptional state in these tumors is therefore only loosely constrained by lineage.

Four clones were represented by at least 20 cells in both a primary tumor and a matched metastasis (Fig. 6g). In each of these, the dominant program was retained at both sites, with differences limited to shifts in the relative use of other programs. The same was true in 103T2, where 4 of 5 paired lineages retained their dominant program across sites. Together with the preceding results, this indicates that transcriptional state in these tumors could be stably propagated within a lineage once established, including to its metastatic derivatives, but is not determined by lineage identity itself, as shown by the divergence of the same founder lineage in two different hosts.

Taken together, the two xenograft models establish that clonal selection shapes MRT growth *in vivo*, with each animal dominated by a largely distinct set of lineages despite receiving the same starting population. The transcriptional programs themselves, by contrast, recurred: lineages that expanded independently, in separate hosts, converged on the same restricted programs, and the one lineage that dominated in two animals adopted a distinct program composition in each. Metastasis arose from lineages already expanded in the matched primary tumor, and no program marked the metastatic compartment or any anatomical site. Many of the lineages that established a primary tumor therefore contributed to metastatic disease, rather than a rare subpopulation in a specialized state. Lineage identity, then, shapes transcriptional state without determining it.

## Discussion

Here, we combined lineage barcoding with single-cell transcriptomics to follow clonal and transcriptional dynamics in two patient-derived models of malignant rhabdoid tumor of the kidney, during long-term culture and grown as orthotopic xenografts. We show, for the first time, clonal dynamics in the context of MRT, a tumor driven by derailed epigenetic gene regulation rather than mutation accumulation caused by genetic instability. Four observations recur across models. First, clonal complexity collapses rapidly and reproducibly, both during passaging and at engraftment. Second, which lineages come to dominate *in vivo* is largely unpredictable and differs between animals given the same starting population. Third, transcriptional programs behave differently from clones: the same states recur across lineages that expanded independently, including in separate hosts. Fourth, that recurrence is bounded by the tumor of origin, with the two models sharing their least differentiated states.

The most severe clonal restriction we observed occurred at engraftment, where barcode complexity fell by orders of magnitude within the first few weeks. That evenness was essentially unchanged over the same interval argues that this loss was a sampling event rather than a selective one: random loss of lineages reduces their number without altering the distribution of clone sizes, whereas preferential outgrowth would be expected to skew it. Comparable clonal restriction has been reported in xenografts of established cancer cell lines, where a small minority of injected cells accounts for nearly all recovered progeny, an effect described as clonal dominance^16^. Our data indicate that in this system the bottleneck largely precedes selection *in vivo*, and that the differential expansion we observe subsequently acts on the small pool of lineages that survive it.

Following engraftment, the two models diverged in how clonal composition evolved during tumor progression. In 103T2, complexity plateaued while evenness continued to fall, indicating expansion within a persisting pool of lineages. In 60T2, however, lineages continued to be lost until tumors were effectively monoclonal. The identity of the winners was correspondingly less reproducible in 103T2, where dominant clones were essentially private to each animal. In 60T2, early tumors shared much of their abundant repertoire, but these shared lineages were largely those already most abundant in the injected pool, reflecting founder representation rather than convergent selection. Taken together, this argues against strong deterministic selection on heritable, clone-intrinsic fitness differences: if such differences dominated, the same lineages should have prevailed across animals receiving the same starting population^25,38,39^. That the lineages favored in culture were largely distinct from those favored *in vivo* reinforces this, and indicates that the two environments impose different constraints.

Transcriptional programs, by contrast, were strikingly recurrent. In both models and in both settings, clustering lineages by program composition grouped clones that had expanded independently, in separate cultures, and in separate hosts, indicating convergence on a limited repertoire of states. This is consistent with the broader observation that malignant cells occupy recurrent transcriptional meta-programs comprising both generic programs, such as cell cycle and stress, and programs specific to a tumor type or lineage^40^. Our data map onto this distinction directly: the programs shared between our two models and modalities were a proliferative program and broadly used progenitor-like states, whereas the differentiated programs were specific to a model or a setting.

The divergence between the two lines is consistent with their developmental origins. MRTs arise from the neural crest, arrested en route to mesenchymal differentiation, and patient origin accounts for most of the transcriptional variation among MRT organoid cells^36^. 103T2 and 60T2 occupy distinct positions along this trajectory, and the programs specific to one line or the other are those these positions would predict. A block at this branch point leaves a wide range of fates open, and different tumors need not resolve toward the same ones. Our data support this: even where the two lines yielded programs with similar functional annotations, they did so through largely distinct genes. Given the genomic stability of MRTs^37^, this divergence is unlikely to reflect differences in cooperating driver mutations. *SMARCB1* loss has instead been shown to activate distal enhancers in a patient-specific manner^41^, providing a mechanism by which the same lesion can yield different transcriptional output in different tumors.

The relationship between lineage and transcriptional state was more nuanced than either full determination or independence. Programs differed markedly in how clone-restricted they were, with generic states used broadly and differentiated states confined to a minority of lineages. In most paired comparisons, a lineage also maintained its dominant program between a primary tumor and its metastases. Yet the one lineage that became dominant in two independent hosts adopted program compositions no more similar to each other than to unrelated clones. Together, these findings suggest that transcriptional states can remain stable within a lineage without their precise composition being rigidly specified by the founder cell. Lineages were also able to change state: in the organoid cultures, individual lineages tracked from early to late passage shifted their program composition substantially, and one xenograft lineage was dominated by different programs in a primary tumor and its metastasis. Part of the transcriptional reorganization we observe therefore reflects plasticity within persisting lineages rather than the replacement of one set of clones by another. A similar distinction has been drawn in other systems, where heritable single-cell expression programs persist over multiple divisions without being genetically encoded^42^, and where the proliferative fate of drug-tolerant persisters is lineage-dependent in the absence of genetic differences^21^. Stability within a lineage was not absolute in our models, however: one lineage was dominated by different programs in a primary tumor and its metastasis, and in the organoid cultures individual lineages shifted composition substantially between early and late passage, indicating that the conditions under which a lineage retains its state are not uniform.

For metastasis, the barcode data were largely consistent across both models: metastases were generally dominated by lineages that had already expanded in the matched primary tumor, rather than by rare subpopulations. Metastases showed a further reduction in clonal diversity relative to their primaries, consistent with an additional bottleneck at dissemination. Barcoding of triple-negative breast cancer patient-derived xenografts likewise found that metastatic clones were traceable to clones detected in the matched primary tumor. However, primary-tumor clone abundance did not predict contribution to metastatic disease, with relatively minor primary clones accounting for much of the metastatic burden^20^. The 103_4 metastatic lineage provided an example of such disproportionate metastatic representation in our models, although this pattern was the exception rather than the rule.

Despite clonal restriction, metastatic dissemination was not associated with a characteristic transcriptional state. Program composition showed no correspondence with anatomical site: two metastases at different sites within one animal could be near-identical, while two metastases from the same organ in another animal differed substantially. In the one lineage whose program composition differed markedly between sites, the program dominating the metastasis was already present in a minority of cells in the primary tumor, leaving open whether the metastasis arose from those cells or acquired the state independently.

### Limitations

The conclusions of this study are subject to several limitations. First, only two patient-derived organoid lines were examined, and while their contrasting behavior is informative, it does not establish whether either is more representative of MRTK generally. Second, our view of transcriptional state *in vivo* is restricted to a single point in time. Single-cell profiling was performed only on established tumors at the experimental endpoint, so we cannot say when the programs we describe were established, or whether the transcriptional composition of a tumor changes as it grows in the way that its clonal composition does. The bulk barcode time course addresses clonal dynamics across tumor development, but is itself cross-sectional: because tumors were harvested at sacrifice, trajectories are inferred from comparisons between animals rather than followed within them. Finally, the number of cells recovered varied by orders of magnitude between samples, and metastases in particular were often small. This restricted the number of paired clone-level comparisons that could be made, and means that the absence of a detectable difference in the smaller samples may reflect limited power.

## Conclusion

Together, these findings provide the first demonstration that clonal evolution shapes MRTK growth. Clonal restriction was substantial in every culture and every animal, and each tumor came to be dominated by a largely distinct set of lineages. The transcriptional states available to these tumors, by contrast, were drawn from a limited and recurring repertoire, on which expanding lineages independently converged. Metastasis drew on many of the lineages that had expanded in the primary tumor, rather than on a specialized state confined to a few. What recurs across our models and settings is therefore not a particular clone or program, but the capacity of lineages to occupy, and to move between, a constrained set of transcriptional states.

---

**Supplementary Figure S1:**
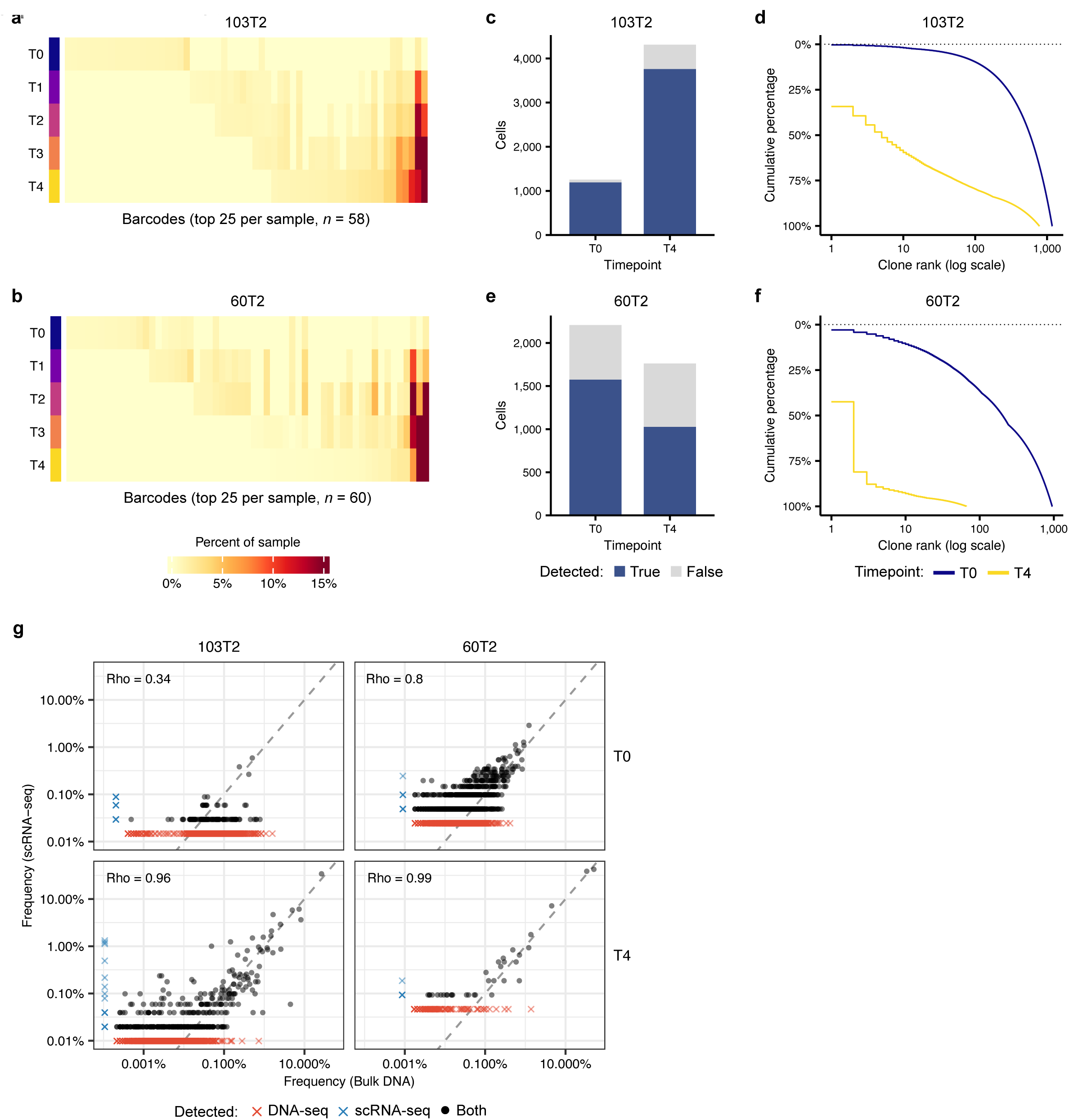
Clonal composition and barcode recovery in MRTK organoid cultures. **(a, b)** Frequency of the union of the 25 most abundant barcodes per sample across the passaging time course for 103T2 (a, *n* = 58 barcodes) and 60T2 (b, *n* = 60 barcodes). Rows are timepoints; columns are barcodes. **(c, e)** Number of cells recovered by scRNA-seq at T0 and T4 in 103T2 (c) and 60T2 (e), divided into cells with and without a detectable barcode. **(d, f)** Cumulative barcode abundance against clone rank (log scale), derived from scRNA-seq, for 103T2 (d) and 60T2 (f). **(g)** Barcode frequencies measured by bulk DNA-seq against those measured by scRNA-seq of the same sample, for each line and timepoint. Dashed line indicates the identity. Barcodes detected by only one assay are plotted at that assay’s detection floor and marked with crosses. Spearman correlations are computed on barcodes detected by both assays.

**Supplementary Figure S2:**
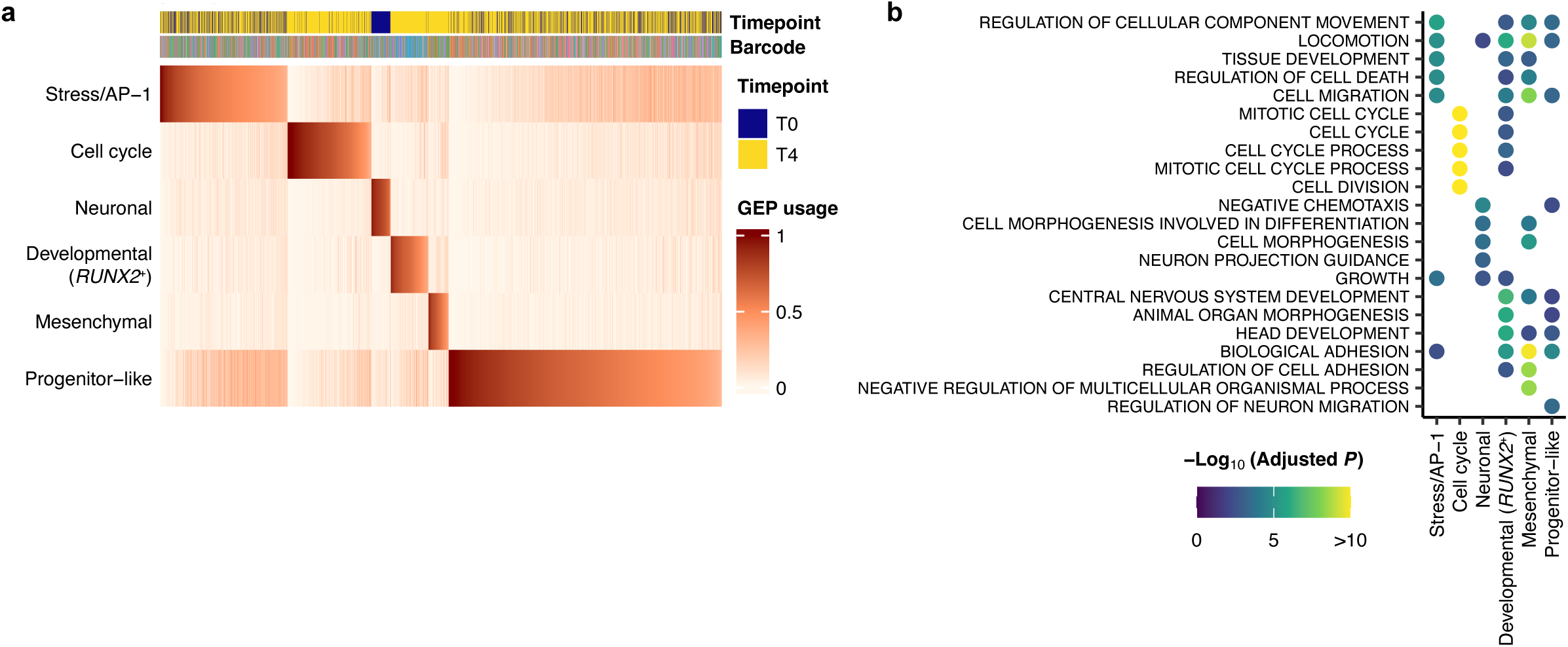
Program usage and functional enrichment in 103T2 organoids. **(a)** Per-cell usage of each of the six 103T2 organoid programs. Columns are cells, ordered by assigned program and by usage of that program; annotation tracks indicate timepoint and barcode identity. **(b)** Top 5 gene ontology terms enriched among the top-weighted genes of each program. Point color indicates the adjusted *P* value on a −log₁₀ scale; terms with no significant enrichment for a given program are left blank.

**Supplementary Figure S3:**
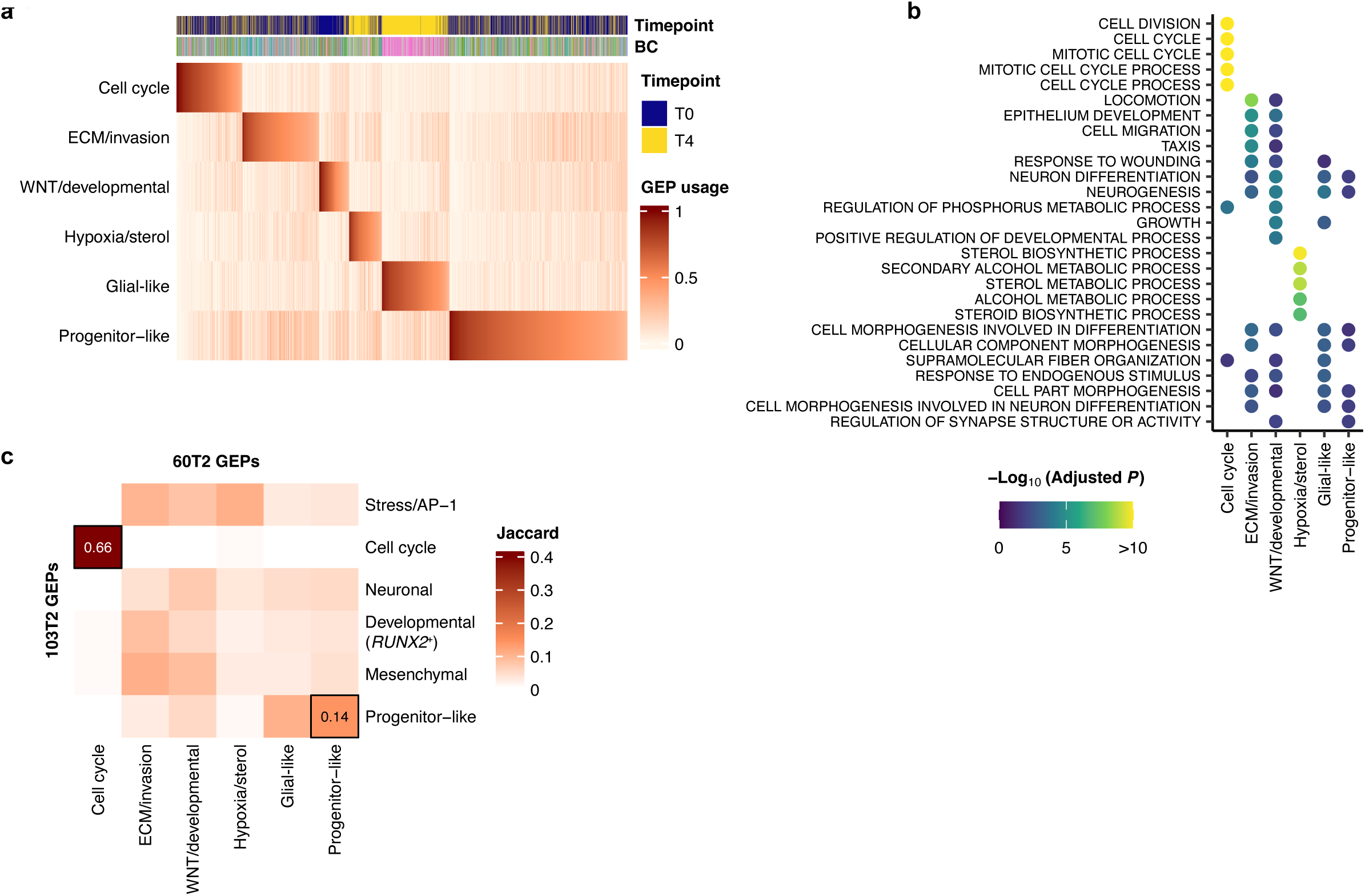
Program usage, functional enrichment and cross-line comparison for 60T2 organoids. **(a)** Per-cell usage of each of the six 60T2 organoid programs. Columns are cells, ordered by assigned program and by usage of that program; annotation tracks indicate timepoint and barcode identity. **(b)** Top 5 gene ontology terms enriched among the top-weighted genes of each program. Point color indicates the adjusted *P* value on a −log₁₀ scale; terms with no significant enrichment for a given program are left blank. **(c)** Jaccard index between the top 100 genes of each 103T2 program (rows) and each 60T2 program (columns). Boxed cells and values indicate pairs exceeding the 99th percentile of a null distribution generated by permuting gene labels.

**Supplementary Figure S4:**
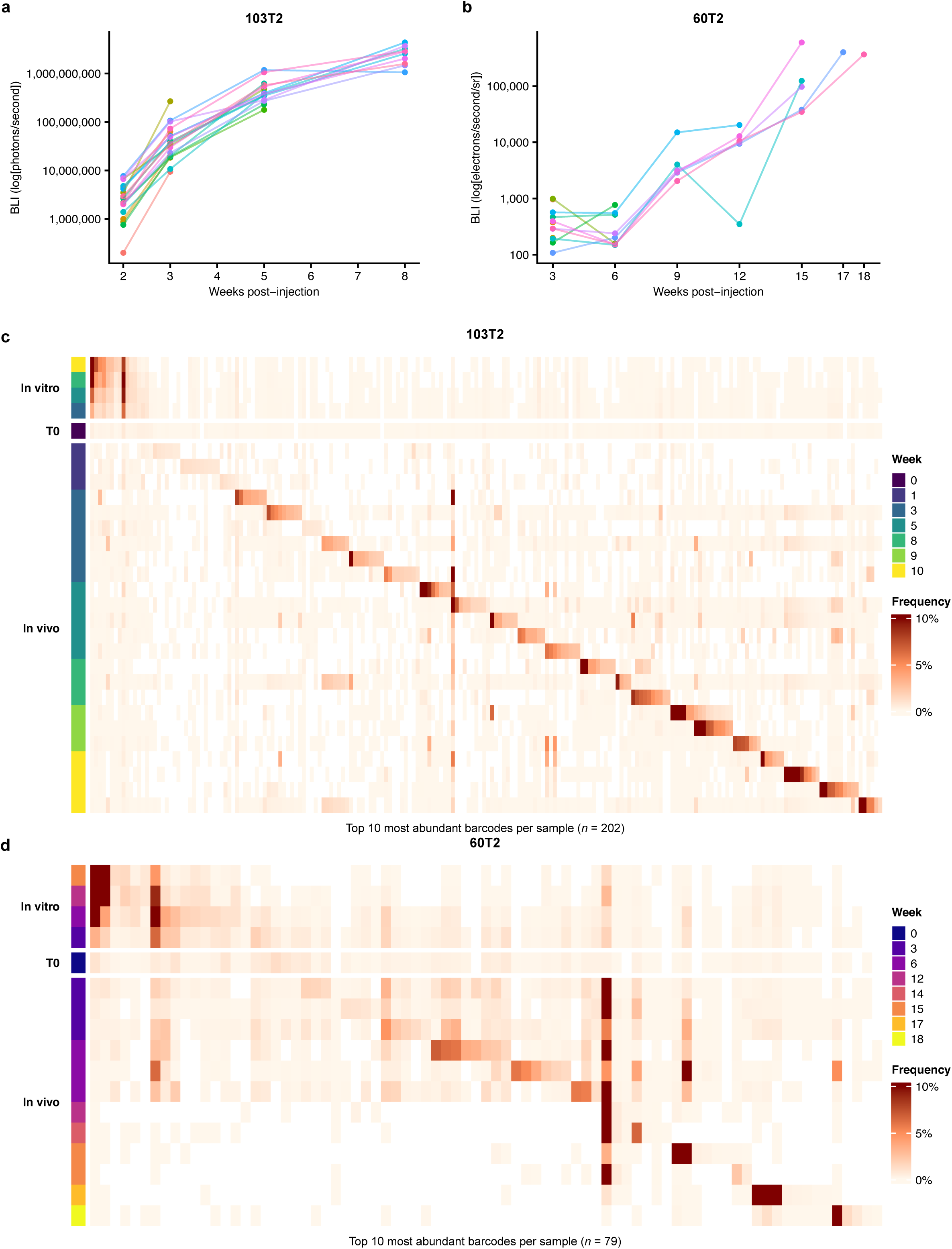
Xenograft growth kinetics, and comparison of clonal composition between *in vitro* and *in vivo* settings. **(a, b)** Bioluminescence imaging (BLI) of primary xenograft tumors derived from 103T2 **(a)** and 60T2 **(b)** organoids. Each line represents one animal, measured at the indicated weeks post-injection; measurements end at the timepoint at which each mouse was sacrificed. **(c, d)** Frequency of the union of the ten most abundant barcodes per sample, combining serially passaged organoids, the injected organoid pool (T0), and xenograft tumors, for 103T2 (a, *n* = 202 barcodes) and 60T2 (b, *n* = 79 barcodes). Rows are samples, grouped by modality and ordered by week collected; columns are barcodes, ordered by the sample in which each reaches its highest frequency.

**Supplementary Figure S5:**
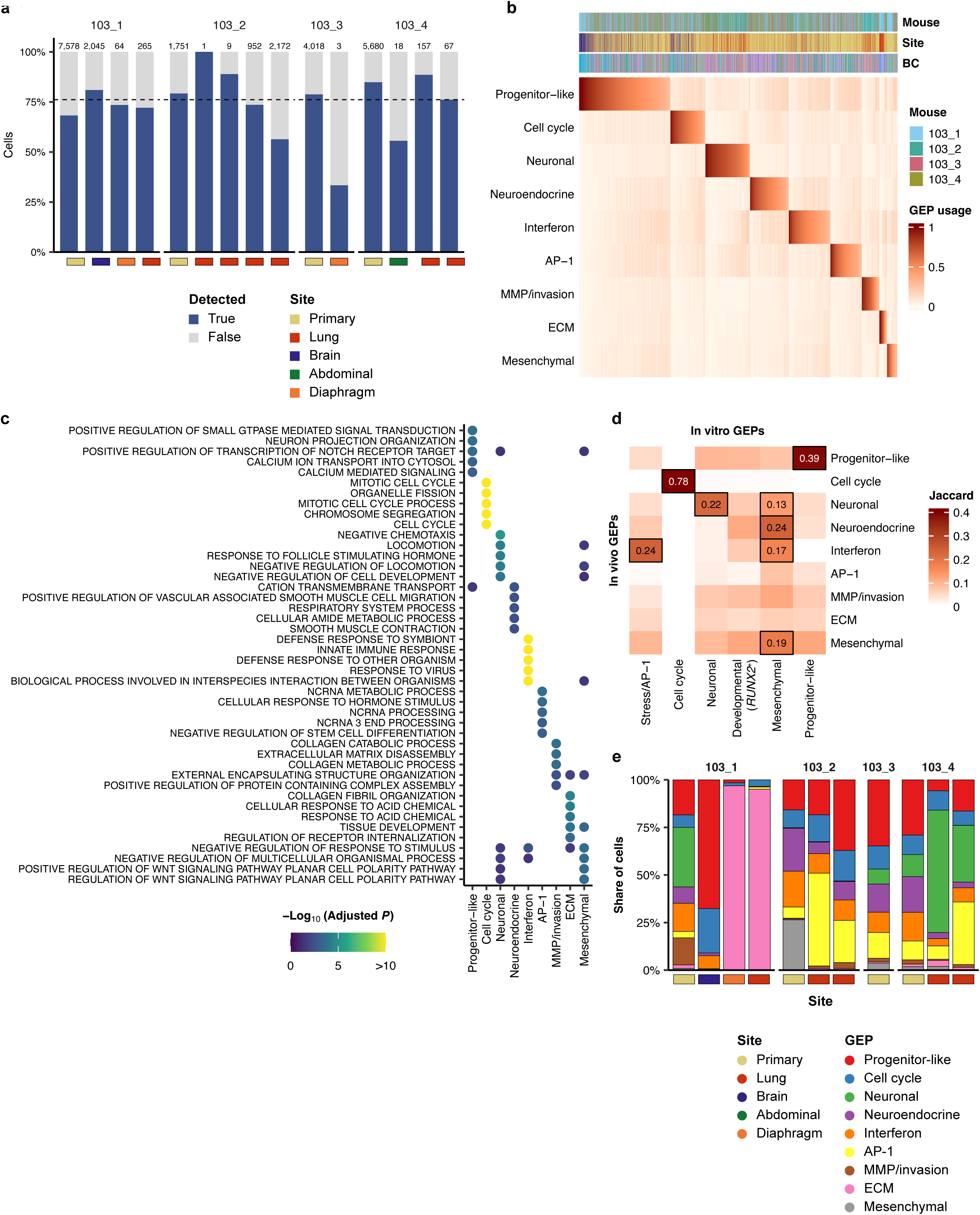
Barcode recovery, program usage and functional annotation for 103T2 xenograft tumors. **(a)** Proportion of cells with and without a detectable barcode in each 103T2 *in vivo* sample, grouped by mouse. The total number of cells per sample is given above each bar and the anatomical site is indicated below. Dashed line, median detection rate across samples. **(b)** Per-cell usage of each of the nine *in vivo* programs. Columns are cells, ordered by assigned program and by usage of that program; annotation tracks indicate mouse, site and barcode identity. **(c)** Top 5 gene ontology terms enriched among the top-weighted genes of each program, as in Supplementary Figure S2b. **(d)** Jaccard index between the top 100 genes of each 103T2 *in vivo* program (rows) and each 103T2 organoid program (columns). Boxed cells and values indicate pairs exceeding the 99th percentile of the permutation null. **(e)** Proportion of cells assigned to each program in each sample, grouped by mouse. The strip below each bar indicates anatomical site.

**Supplementary Figure S6:**
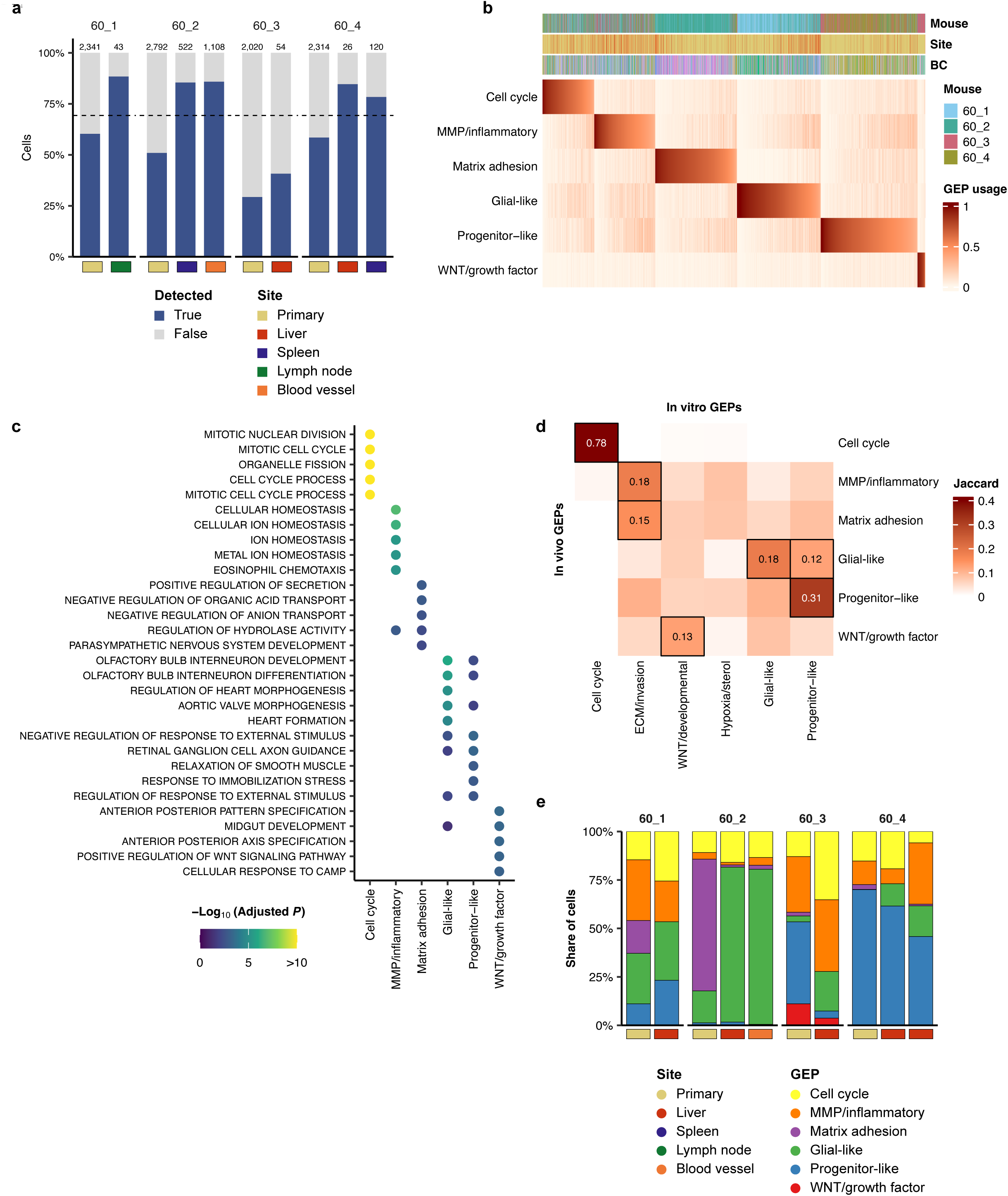
Barcode recovery, program usage and functional annotation for 60T2 xenograft tumors. **(a)** Proportion of cells with and without a detectable barcode in each 60T2 *in vivo* sample, grouped by mouse. The total number of cells per sample is given above each bar and the anatomical site is indicated below. Dashed line, median detection rate across samples. **(b)** Per-cell usage of each of the six *in vivo* programs. Columns are cells, ordered by assigned program and by usage of that program; annotation tracks indicate mouse, site and barcode identity. **(c)** Top 5 gene ontology terms enriched among the top-weighted genes of each program, as in Supplementary Figure S2b. **(d)** Jaccard index between the top 100 genes of each 60T2 *in vivo* program (rows) and each 60T2 organoid program (columns). Boxed cells and values indicate pairs exceeding the 99th percentile of the permutation null. **(e)** Proportion of cells assigned to each program in each sample, grouped by mouse. The strip below each bar indicates anatomical site.

**Supplementary Table S1**: Summary of mouse cohorts and experimental timepoints *in vivo*. Number of mice euthanized at specific weeks post-injection, either at planned timepoints or after reaching a humane endpoint.

**Supplementary Table S2**: Primers and PCR settings for NGS library preparation. List of PCR 1 and PCR 2 primers used for NGS library preparation and PCR cycling steps.

## Methods

### Lentiviral barcode library construction

To generate the lentiviral barcode library, a semi-random barcode cassette was synthesized and cloned into the lentiviral backbone pLV-EF1α-Puro-P2A-H2B-mNeonGreen-polyA (VectorBuilder)^35^. This vector contains an EF1α promoter driving a single open reading frame encoding puromycin resistance and a nuclear-localized fluorescent reporter (H2B-mNeonGreen). The barcode cassette was inserted into the 3′-UTR of mNeonGreen between the unique AscI and NsiI restriction sites, approximately 170 bp upstream of the bGH polyadenylation signal to facilitate barcode detection during RNA sequencing. The barcode consisted of four repeats of the semi-random motif ABNNNB followed by a terminal A ((ABNNNB)₄A), where N represents any nucleotide and B represents any nucleotide except adenine. This design yields a theoretical barcode diversity of approximately 10¹¹ unique sequences. Oligonucleotides containing the barcode region flanked by M13 priming sites and AscI/NsiI restriction sites were synthesized by Integrated DNA Technologies (IDT).

Double-stranded barcode fragments were generated by primer extension using a 22 bp forward primer and an 81 bp reverse primer containing the barcode sequence. Primers were denatured at 98 °C followed by slow cooling to room temperature. dNTPs and Klenow fragment polymerase (Promega, M2201) were then added, and the reactions were incubated for 1 h at 37°C. Five parallel primer extension reactions were performed and subsequently pooled to increase DNA yield. DNA was ethanol precipitated using sodium acetate and GlycoBlue, washed with 70% ethanol, and resuspended in TE buffer.

The barcode fragments were amplified using primers containing Gibson assembly overhangs and Phusion Hot Start HF polymerase (NEB) and purified using the QIAquick PCR purification kit (Qiagen). The pLV-EF1a-Puro-P2A-H2B-mNeonGreen-polyA backbone was digested with AscI (FastDigest SgsI, ThermoFisher) and NsiI (FastDigest Mph1103I), followed by XbaI (NEB) and KpnI-HF (NEB) digestion to remove residual undigested plasmid and treated with TSAP phosphatase (Promega) to prevent self-ligation. Barcode inserts were cloned into the prepared backbone using Gibson assembly with a 100-fold molar excess of insert. Multiple parallel reactions were pooled and transformed into NEB Stable competent *E. coli* (C3040H) following the manufacturer’s high-efficiency protocol. Transformed bacteria were plated on LB-agar containing ampicillin and incubated at 30°C for approximately 44 h. Colonies were scraped, pooled, expanded in LB medium with ampicillin, and plasmid DNA was extracted using the NucleoBond Xtra Maxi kit (Macherey-Nagel). Virus production was performed as previously described^43^.

### Patient-derived organoids: cell culture and transduction

Experiments with human material were approved by the medical ethical committee of the Erasmus Medical Center (Rotterdam, the Netherlands) and Princess Máxima Center for Pediatric Oncology (Utrecht, the Netherlands). Informed consent was obtained from the parents of all participants. Approval for use of the subject’s tissue samples within the context of this study has been granted by the Máxima biobank and data access committee (https://research.prinsesmaximacentrum.nl/en/core-facilities/biobank-clinicalresearch-committee) (biobank request nr. PMCLAB2018.005 and PMCLAB2018.006).

MRTK organoids were established previously and cultured as described^32^. Briefly, organoids were seeded in growth factor-reduced BME (R&D Systems) and cultured in kidney organoid medium (KOM). KOM consists of AdDF+++ (Advanced DMEM/ F12 (Gibco) containing 1X Glutamax (Thermo Fisher Scientific), 10 mM HEPES (Thermo Fisher Scientific) and 1X Penicillin-Streptomycin (Merck Millipore)), supplemented with 1.5% B27 supplement (Gibco), 10% R-spondin-conditioned medium, EGF (50 ng/ml, Peprotech), FGF-10 (100 ng/ml, Peprotech), N-acetylcysteine (1.25 mM, Sigma), Rho-kinase inhibitor Y-27632 (10 μM, Abmole) and A83-01 (5 μM, Tocris Bioscience). Medium was changed every 3-4 days and organoids were passaged weekly by mechanical dissociation.

MRTK organoids were transduced^43–45^ with pLKO.1-Ubc-luciferase-blast^46^ to follow tumor progression *in vivo* using bioluminescence imaging (BLI). Organoids were selected with blasticidin (10 μg/mL). 1,000,000 organoids stably expressing luciferase were subsequently transduced with the lentiviral barcode library, using 30 μL or 10 μL of virus for 103T2 model or 60T2 model respectively. 72h after transduction, the percentage of mNeon+ organoids was quantified by FACS to estimate the infection efficiency. Based on the percentage of mNeon+ cells (74% for 103T2 organoids and 40% for 60T2 organoids), we estimated an MOI of 1.35 and 0.5 respectively (calculated using a Poisson distribution model^25^).

Immediately after, organoids were put under puromycin selection (1 μg/mL) and passaged for 3 weeks before orthotopic transplantation, to remove any remaining virus. Organoid were kept in culture under mild antibiotic selection (puromycin 0,25 μg/mL, blasticidin 2,5 μg/mL) throughout the experiment to ensure that both constructs were maintained. Excess organoids from each passage were viably frozen for NGS and scRNA-seq analyses.

### Animal studies

All procedures were performed in accordance with the Dutch Experiments on Animals Act 2014 and approved by the Dutch Competent Authority (AVD39900202216405) and the Animal Welfare Body of the Princess Máxima Center (PMC.65.3067.1901.1, PmcDro16405.04).

Female NOD.Cg-*Prkdc^scid^ Il2rg^tm1Wjl^*/SzJ (NSG) mice, aged 7–14 weeks, were used as acceptors for orthotopic transplantation. Animals were kept in individually ventilated cages (Innovive or Tecniplast, Greenline) with Corn Cob bedding, dome, nesting sheet and ladder (Innovive, USA) at a temperature of 20.5°C to 23.5°C, humidity of 45% to 65%, 50 air exchanges per hour in the cages, and a 14/10-hour light/dark cycle with the lights on at 6 AM. Mice were fed a standardized mouse diet (RM3; SDS, Germany) and provided drinking water (acidified) ad libitum. All materials, including individually ventilated cages, lids, feeders, bottles, bedding, and water were autoclaved before use. Mice were housed under specific (and opportunist) pathogen-free conditions (Replace monitoring; IDEXX, Germany).

The day before transplantation, organoids were collected, washed in cold AdDF+++ and incubated for 5 min at 37°C with TrypLE Express (Invitrogen, 1260510) containing 10 μM Y-27632 to remove BME. Approximately 250,000 cells were replated in 10 μL BME droplets and allowed to recover overnight at 37 °C and 5% CO₂.

Renal capsule transplantation protocol was adapted from previous studies^46^. Recipient mice were anesthetized with isoflurane. Through a flank incision, the left kidney was exposed and a small opening was created in the renal capsule at the lower pole of the kidney. A single BME droplet containing 250,000 organoids was mixed with 1uL Fast green FCF dye (Sigma) and gently injected beneath the renal capsule using a PE50 catheter, pre-coated with Sigmacote, attached to a Hamilton syringe.

Tumor growth was monitored through bioluminescence imaging (BLI) every 3 weeks, starting from 21 days after transplantation. Mice received a subcutaneous injection of D-luciferin (150 mg/kg, Revvity), were anesthetized with isoflurane and placed in a dark chamber. After a 12 minute incubation period, BLI signal was detected using IVIS or MILabs Optical Imaging System (60T2). When animals reached a humane endpoint, an additional BLI measurement was performed under terminal anesthesia.

Mice were euthanized by cervical dislocation under terminal anasthesia. The left kidney and distant organs (liver, lungs, diaphragm, brain, spleen and lymph nodes) were collected and inspected under an epifluorescent microscope (Leica, M205 FA). Primary tumors and metastatic lesions identified by H2B-mNeon fluorescence were then microdissected and viably frozen. Larger tissues were minced into ∼1 mm³ pieces and thoroughly mixed to randomize barcode distribution before viable freezing.

### gDNA extraction, NGS library preparation and sequencing

gDNA was extracted from organoids or tissues using ReliaPrep™ gDNA Tissue Miniprep System (Promega, A2051). When sufficient tissue was available, three independent replicates of 25 mg were processed to maximize barcode recovery. Tissues were homogenized with a Dounce homogenizer before incubation with Proteinase K.

Lineage tracing barcodes were amplified with a two-step PCR protocol. For PCR 1, a total of 1.65 mg of gDNA extracted from organoids (corresponding to approximately 250,000 cells) and 5 mg of gDNA extracted from tissues were used as template. The DNA was distributed across parallel reactions, with a maximum of 500 ng of DNA per reaction and a total reaction volume of 50 µL per tube. PCR 1 was performed using Phusion™ Hot Start II DNA Polymerases (ThermoFisher, F549L) and primers designed to flank the lineage tracing barcode, with the forward primer annealing to the mNeon sequence and the reverse primer to the poly(A) sequence.

After PCR 1, amplicons from parallel reactions were pooled and cleaned up with AMPure XP Beads (BeckMan Coulter) and excess primers were digested using ExoSap-IT (ThermoFisher). DNA concentration was quantified with Qubit and the expected amplicon size (349 bp) was confirmed using the Agilent Bioanalyzer High Sensitivity DNA assay.

1 ng of purified DNA from PCR 1 was used as template for PCR 2. A single reaction of 20 µL volume was set up for each sample. PCR 2 was performed using Phusion™ Hot Start II DNA Polymerases and primers containing the following elements: a sequence complementary to PCR 1 primers, a 10-bp Nextera unique dual index for multiplexing and the i5/i7 Illumina adapter sequences. PCR amplicons were purified with Zymo DNA Clean&Concentrator kit. DNA concentration was quantified with Qubit and the expected amplicon size (422 bp) was confirmed using the Agilent Bioanalyzer High Sensitivity DNA assay. Libraries were diluted to 4 nM, pooled and sequenced on Illumina NextSeq2000 with a target depth of approximately 1 million reads per sample. When sequencing runs contained only these libraries, 50% PhiX was added to increase sequence diversity. Primers and PCR cycling conditions are listed in Supplementary Table S2.

### Analysis of NGS data

Barcodes extraction strategy was adapted from previous studies^47^ using functions from the genBaRcode package in R^48^. FASTQ files were demultiplexed and quality-filtered to retain reads with a mean Phred score > 30. From the filtered FastQ files, 300,000 reads per sample were subsampled to equalize sequencing depth across samples. Barcode sequences were extracted from the subsampled reads by matching the backbone sequence “ABNNNBABNNNBABNNNBABNNNBA”. Barcode counts from all samples were then combined into a single matrix. To build the barcode reference list, barcodes with a total read count > 10 across all samples were selected and merged using a Hamming distance (HD) threshold = 2. Barcodes with total read counts between 3 and 10 were then added to this high-confidence set, and the expanded barcode set was subjected to a second round of HD = 2 correction. Finally, the barcode reference list was intersected with the combined barcode count matrix, and shared barcodes were retained for downstream analyses.

Barcode counts were converted to relative abundances within each sample by dividing by the sample total. Clonal complexity was quantified as the number of unique barcodes detected per sample, counting barcodes supported by at least 5 reads after downsampling. Clonal evenness was quantified using Pielou’s evenness index, J = H′/ln(S), where H′ is the Shannon index, H′ = −Σ pᵢ ln pᵢ, computed over the relative abundances pᵢ of the S barcodes detected in that sample (implemented in the vegan R package^49^, v2.7-1). Cumulative abundance curves were constructed by ranking barcodes within each sample by decreasing relative abundance and plotting the cumulative sum against rank. The number of clones accounting for half of all reads was taken as the smallest number of top-ranked barcodes whose cumulative abundance reached 50%.

### Sample processing for scRNA-sequencing

Viably frozen organoids and tissues were thawed in a 37 ᵒC water bath and transferred to a 15 mL Falcon tube. Cold AdDF+++ was immediately added dropwise and samples were centrifuged at 300 x *g* for 5 min at 4 ᵒC. Primary tumors were further minced using scalpels prior to enzymatic dissociation, whereas metastases were directly dissociated. Tissue samples were enzymatically dissociated with Collagenase II brown powder (Thermo Fisher) dissolved in AdDF+++ supplemented with RhoKi 10 μM. Collagenase II was used at 5 mg/mL for primary tumors and 2,5 mg/mL for metastases. Samples were incubated in a shaking incubator at 37 ᵒC and 200 rpm for approximately 45 min and subsequently washed with AdDF+++. Organoids were mechanically dissociated as previously described^32^. Following dissociation, organoid and tissue samples were resuspended in FACS buffer, filtered through a 70 µM filter and stained with DAPI and Draq5^2^. Viable mNeon+ cells (DAPI-, Draq5+) were FACS sorted using a Sony Cell sorter. Following sorting, cells were counted manually using a Bürkett chamber and cell viability was estimated using Trypan Blue solution.

Single-cell gene expression libraries were prepared using the Chromium platform (10x Genomics). Libraries from 60T2 samples were multiplexed using on-chip multiplexing (OCM). 103T2 samples were either labelled with Cell Multiplexing Oligos (CMOs) prior to FACS sorting (primary tumors and organoids at T4) and demultiplexed, or processed as independent libraries (metastases and organoids at T0). Libraries were sequenced on NovaSeq 6000 (Illumina).

### Analysis of scRNA-seq data

#### Read alignment, demultiplexing and quantification

Raw sequencing reads were processed with Cell Ranger (10x Genomics, version 7.1.0) and mapped against the GRCh38 human reference transcriptome. To allow detection of the lineage tracing barcodes, the reference was supplemented with an additional contig corresponding to the barcoded transgene (EF1α–Puro–P2A–H2B–mNeonGreen–polyA, with the barcode region represented as “ABNNNBABNNNBABNNNBABNNNBA”. Reads carrying the lineage barcode therefore aligned to this contig and were recovered from the Cell Ranger BAM output for barcode extraction (below). 103T2 primary tumors (excluding mouse 103_2) and T4 organoids, which were labelled with Cell Multiplexing Oligos (CMO), were demultiplexed on the basis of CMO counts using cellranger ocm with the assignment defaults; cells called as multiplets or unassigned were discarded. 60T2 libraries, prepared with on-chip multiplexing (OCM), were demultiplexed using the cellranger multi OCM workflow. 103T2 metastases and T0 organoids were processed as independent libraries.

#### Quality control and filtering

Count matrices were analyzed in R (version 4.4.3) using Seurat (version 5.1.0)^50^. Doublets were identified using scDblFinder (v1.18.0)^51^ after having first removed cells with fewer than 250 UMIs. Cells were excluded if they were called doublets in at least 9 out of 10 independent runs of scDblFinder, setting “dbr.sd = 1”. Singlet cells were then retained if they expressed at least 2000 genes, had fewer than 50,000 UMIs, and had a mitochondrial read fraction below 15%.

#### Normalization, dimensionality reduction and visualization

Within each subset of the data (103T2 Organoids, 60T2 Organoids, 103T2 Xenografts, and 60T2 Xenografts), UMI counts were log-normalized with a scaling factor of 10,000, the 2000 most variable genes were selected, and the data were scaled and centered, regressing out the effect of cell cycle phase. Principal component analysis (PCA) was then performed, excluding mitochondrial and ribosomal protein-encoding genes as well as the barcode construct. Batch correction was performed on the resulting principal components using Harmony^52^, with either the timepoint (organoid data) or mouse (xenograft data) used as the group variable. The first 30 batch-corrected principal components were then used to construct UMAP embeddings for visualization.

#### Lineage barcode recovery from single-cell data

Lineage barcodes were recovered in R from the reads aligning to the barcode construct contig. The 25-nt lineage barcode was extracted from each read sequence by matching the (ABNNNB)₄A motif (allowing zero mismatches) and associating each barcode with its corresponding 10x cell barcode. Reads were then collapsed to unique UMIs per cell-lineage barcode combinations. Lineage barcodes recovered in this way were filtered to retain only those found in the reference barcode list generated from the bulk DNA-seq data (see “Analysis of NGS data”). Where more than one lineage barcode was identified in a single cell, the barcodes were appended and the cell was flagged. Cells without a detectable lineage barcode were retained for all transcriptome-level analyses (UMAP embedding, NMF decomposition) but excluded from clone-level analyses.

#### Comparison of clonal composition between bulk DNA-seq and scRNA-seq

For each sample profiled by both assays, barcode frequencies were computed as the fraction of reads (bulk DNA-seq) or the fraction of barcoded cells (scRNA-seq) assigned to each barcode. Spearman correlations were calculated on the set of barcodes detected by both assays; barcodes detected by only one assay were plotted at that assay’s detection floor and excluded from the correlation.

#### Non-negative matrix factorization (NMF)

Gene expression programs (GEPs) were identified using the RcppML^53^ (version 0.3.7.1) implementation of NMF. Within each line, organoid decompositions were performed independently for each timepoint, and independently per mouse for the xenograft data. Input features were selected using the “SelectIntegrationFeatures” function that identified the top 2000 shared variable features across groups for each decomposition. These were again filtered to exclude mitochondrial and ribosomal protein-encoding genes as well as the barcode construct. The count data were then re-normalized using SCTransform^54^, and the centered residual values were extracted for the input features, setting negative values to zero. NMF was then run for ranks k = 3 to 20, with 20 replicates per rank. The final rank was selected by inspecting the stability and reconstruction-error profile across ranks.

Programs were then compared between groups by the Jaccard index of their top 100 weighted genes and clustered by hierarchical clustering of 1 − Jaccard with average linkage, and clusters were defined by cutting the tree at a height of 0.85 (corresponding to a mean Jaccard index of 0.15 between cluster members). Clusters containing programs from at least two groups were taken as recurrent metaprograms, and their gene weights defined as the mean of the constituent programs’ weight vectors. Programs not assigned to a recurrent cluster were retained individually with their original weights. The combined gene by program matrix was then scaled so that each column had unit Euclidean norm. Per-cell program usages were obtained by projecting each cell onto the combined program matrix and normalized to sum to one across programs within each cell. Each cell was additionally assigned to the single program with the highest usage for compositional analyses. Programs were annotated manually from their top-weighted genes together with the gene ontology enrichment described below; annotations are descriptive labels and are not intended to imply exclusive activity of the named process.

#### Gene ontology term enrichment

The top 100 weighted genes of each program were tested for enrichment of Gene Ontology Biological Process terms using a hypergeometric test, implemented in the “fora” function provided by the fgsea R package (1.30.0). P values were adjusted by the Benjamini-Hochberg method.

#### Comparison of programs between lines and between modalities

Programs from different decompositions were compared by the Jaccard index of their top 100 weighted genes. To assess significance, a null distribution was generated by permuting gene labels 1000 times and recomputing all pairwise Jaccard indices; pairs exceeding the 99th percentile of this null were considered to correspond.

#### Clone-level analyses of program usage

Clone-level analyses were restricted to lineages with sufficient cells: at least 10 cells for the organoid datasets and at least 20 cells for the xenograft datasets. Samples contributing fewer than 20 barcoded cells were excluded from the xenograft clone-level analyses.

For each lineage, the fraction of its cells assigned to each program was computed. Enrichment of a program within a lineage was tested against the remaining cells of the same sample using a hypergeometric test (one-sided, testing for over-representation), and P values were adjusted across all lineage-program combinations within a dataset by the Benjamini-Hochberg method; q < 0.05 was considered significant.

For lineages represented in two conditions, the distribution of cells across assigned programs was compared between conditions using Fisher’s exact test on the program by condition contingency table, with P values obtained by Monte Carlo simulation (simulate.p.value = TRUE, B = 100,000 replicates), followed by Benjamini–Hochberg correction across the lineages tested.

#### Sample- and site-level comparisons of program composition

Program composition of a sample was defined as the fraction of its cells assigned to each program. Comparisons of individual program usage between primary tumors and matched metastases were made descriptively per mouse.

## Statistics and reproducibility

All analyses were performed in R (version 4.4.3). Tests were two-sided unless stated otherwise, and multiple testing was controlled by the Benjamini-Hochberg procedure within each analysis; adjusted values are reported as q. No statistical method was used to predetermine sample size, and no cells or samples were excluded other than by the quality control and minimum-cell criteria described above.

## Data and code availability

Raw and processed sequencing data will be deposited in appropriate repositories and made available upon publication. Analysis code will be made available upon final publication.

## Acknowledgements

We are grateful to patients and parents who agreed to participate in our research. We thank the Princess Máxima Center Single Cell Genomics Facility, the FACS Facility and the Animal Research Facility. We thank Marjolein Kes who helped with organoids collection and tissue processing, and Senna Visser who helped with renal capsule transplantations.

## Author contributions

Conceptualization: A.F., L.K., J. Drost. Data Curation: G.P., J. DeMartino, A.F., D.A. Formal Analysis: G.P., J. DeMartino, A.F., D.A. Funding Acquisition: J. Drost. Investigation: G.P., J. DeMartino, A.F., D.A., S.D., J.K. Methodology: G.P., J. DeMartino, A.F., L.K., T.M., J. Drost. Project Administration: G.P., A.F., J. Drost. Resources: G.P., A.F., J.K. Software: G.P., J. DeMartino. Supervision: A.F., J. Drost. Visualization: G.P., J. DeMartino. Writing (original draft): G.P., J. DeMartino. Writing (review and editing): G.P., J. DeMartino, A.F., J. Drost.

